# Wheat *VRN1*, *FUL2* and *FUL3* play critical and redundant roles in spikelet development and spike determinacy

**DOI:** 10.1101/510388

**Authors:** Chengxia Li, Huiqiong Lin, Andrew Chen, Meiyee Lau, Judy Jernstedt, Jorge Dubcovsky

**Author notes:** These authors contributed equally to this work (Chengxia Li, Huiqiong Lin).

## Abstract

The spikelet is the basic unit of the grass inflorescence. In this study, we show that wheat MADS-box genes *VRN1*, *FUL2* and *FUL3* play critical and redundant roles in spikelet and spike development, and also affect flowering time and plant height. In the *vrn1ful2ful3*-null triple mutant, the inflorescence meristem formed a normal double-ridge structure, but then the lateral meristems generated vegetative tillers subtended by leaves instead of spikelets. These results suggest an essential role of these three genes in the determination of spikelet meristem identity and the suppression of the lower ridge. Inflorescence meristems of *vrn1ful2ful3*-null and *vrn1ful2*-null remained indeterminate and single *vrn1*-null and *ful2*-null mutants showed delayed formation of the terminal spikelet and increased number of spikelets per spike. Moreover, the *ful2*-null mutant showed more florets per spikelet, which together with a higher number of spikelets, resulted in a significant increase in the number of grains per spike in the field. Our results suggest that a better understanding of the mechanisms underlying wheat spikelet and spike development can inform future strategies to improve grain yield in wheat.

**SUMMARY STATEMENT:** The wheat MADS-box proteins VRN1, FUL2 and FUL3 play critical and overlapping roles in the development of spikelets, which are the basic unit of all grass inflorescences.

## INTRODUCTION

The grass family (Poaceae) has approximately 10,000 species, including important food crops such as rice, maize, sorghum, barley, and wheat (Kellogg, 2001). The flowers of these species are organized in a unique and diagnostic structure called spikelet (literally “little spike”), which is a compact inflorescence developing within the larger inflorescence (Malcomber et al., 2006). A spikelet typically has two sterile bracts (called glumes) enclosing one or more florets. Each floret includes a carpel, 3 or 6 stamens and two modified scales (called lodicules), all subtended by two bract-like organs, the palea and the lemma (Preston et al., 2009).

Grass inflorescences have been described as a progressive acquisition of different meristem identities that begins with the transition of the vegetative shoot apical meristem (SAM) to an inflorescence meristem (IM). The IM generates lateral primary branch meristems (PBMs) and secondary branch meristems (SBM) that terminate into spikelet meristems (SMs), which generate glumes and lateral floral meristems (FMs) (McSteen et al., 2000). This model has been a useful phenomenological description but is too rigid to explain some grass branching mutants, so a more flexible model is emerging in which meristem fate is regulated by the genes expressed at discrete signal centers localized adjacent to the meristems (Whipple, 2017).

In wheat, shortening of the inflorescence branches results in spikelets attached directly to the central axis or rachis and the formation of a derived inflorescence, a spike, in which spikelets are arranged alternately in opposite vertical rows (a distichous pattern) (Kellogg et al., 2013). In the initial stage, the IM generates a double-ridge structure in which the lower ridges are suppressed and the upper ridges acquire SM identity and form spikelets. The number of spikelets per spike is determined by the number of lateral meristems formed before the transition of the IM into a SM to form the terminal spikelet. In wheat, the growth of the spike is determinate, but the growth of each spikelet is indeterminate, with each SM initiating a variable number of FMs (Ciaffi et al., 2011). The numbers of spikelets per spike and florets per spikelet determine the maximum number of grains per spike and are important components of wheat grain yield potential.

Studies in Arabidopsis, which has a simpler inflorescence than grasses (Malcomber et al., 2006), have shown that MIKC-type MADS-box transcription factors *APETALA1* (*AP1*), *CAULIFLOWER* (*CAL*) and *FRUITFULL* (*FUL*) are critical in the determination of floral meristem identity. In the triple *ap1calful* mutant, the IM is not able to produce flowers and reiterates the development of leafy shoots (Ferrándiz et al., 2000). In rice, combined loss-of-function mutations in *OsMADS14* (ortholog of *VRN1*) and *OsMADS15* (ortholog of *FUL2*) resulted in inflorescences with leaf-like organs on top of the primary branches (Wu et al., 2017). Simultaneous knockdown of *OsMADS14, OsMADS15* and *OsMAD18* (ortholog of *FUL3*) in a *Ospap2* (a SEPALLATA subfamily MADS-box) mutant background eliminated the formation of primary branches, and resulted in the formation of lateral vegetative tillers subtended by leaves (Kobayashi et al., 2012).

MIKC-type MADS-box proteins have a highly conserved MADS DNA-binding domain, an Intervening (I) domain, a Keratin-like (K) domain, and a C-terminal domain. These proteins bind as dimers to DNA sequences named ‘CArG’ boxes, and organize in tetrameric complexes that can recognize different CArG boxes. The multimeric nature of these complexes generates a large number of combinatorial possibilities with different targets and functions (Honma and Goto, 2001); (Theissen et al., 2016).

The closest homologs to the Arabidopsis MADS-box genes *AP1*, *CAL* and *FUL* in the grass lineage are *VERNALIZATION 1* (*VRN1*), *FUL2* and *FUL3*. A phylogenetic analysis of the proteins encoded by these genes (Fig. S1) indicates that the Arabidopsis and grass proteins have independent sub-functionalization stories (Preston and Kellogg, 2006). In the grass lineage, the *VRN1* and *FUL2* clades are closer to each other than to the *FUL3* clade (Preston and Kellogg, 2006). Mutations causing large truncations in the proteins encoded by the two *VRN1* homeologs in tetraploid wheat delayed heading time, but did not alter spikelet morphology or the ability of flowers to form viable grains (Chen and Dubcovsky, 2012). Since *FUL2* and *FUL3* are the closest paralogs of *VRN1*, we hypothesized that they could have redundant spikelet and floral meristem identity functions.

In this study, we combined loss-of-function mutants for the two homeologs of *VRN1*, *FUL2* and *FUL3* to generate double- and triple-null mutants in the same tetraploid background. Characterization of these mutants revealed that *VRN1*, *FUL2* and *FUL3* have overlapping roles in the regulation of flowering time and stem elongation and, more importantly, that they play critical and redundant roles in spikelet development, suppression of the lower ridge and spike determinacy. Individual *vrn1* and *ful2* mutants showed significant increases in the number of spikelets and grains per spike, suggesting that manipulations of these genes may contribute to increasing wheat grain yield potential.

## RESULTS

### Combination of loss-of-function mutations in *VRN1, FUL2* and *FUL3*

We identified point mutations in the A and B genome homeologs of *FUL2* and *FUL3* in an EMS-mutagenized population of the tetraploid spring wheat variety Kronos (Krasileva et al., 2017; Uauy et al., 2009). We selected mutations that generated premature stop codons or modified splice sites. The proteins encoded by these mutant alleles are predicted to have large deletions or complete truncations of the K and C domains (Fig. S2 and Material and Methods) and, therefore, are most likely not functional. We backcrossed each individual mutant two to three times into a Kronos *vrn-2* null background (Distelfeld et al., 2009b) to reduce background mutations. This genetic background was used to avoid the extremely late flowering of plants carrying the *vrn1*-null mutation in the presence of the functional *VRN2* flowering repressor (Chen and Dubcovsky, 2012). All mutants described in this study are in Kronos *vrn2*-null background, which is referred to as “Control” in all figures.

We intercrossed the A and B genome homeologs for each gene and selected plants homozygous for both mutations. For simplicity, mutants with loss-of-function mutations in both homeologs will be referred to as null mutants (e.g. *vrn1*-null). The *ful2*-null and *ful3*-null mutants were crossed with *vrn1*-null (Chen and Dubcovsky, 2012) to generate *vrn1ful2*-null and *vrn1ful3*-null mutants, which were intercrossed to generate all eight homozygous *VRN1*, *FUL2* and *FUL3* allele combinations including the triple *vrn1ful2ful3*-null mutant. These eight genotypes were analyzed for stem length (Fig. 1A) and number of leaves (Fig. 1B) using three-way factorial ANOVAs (Fig. 1C).

**Fig. 1.**
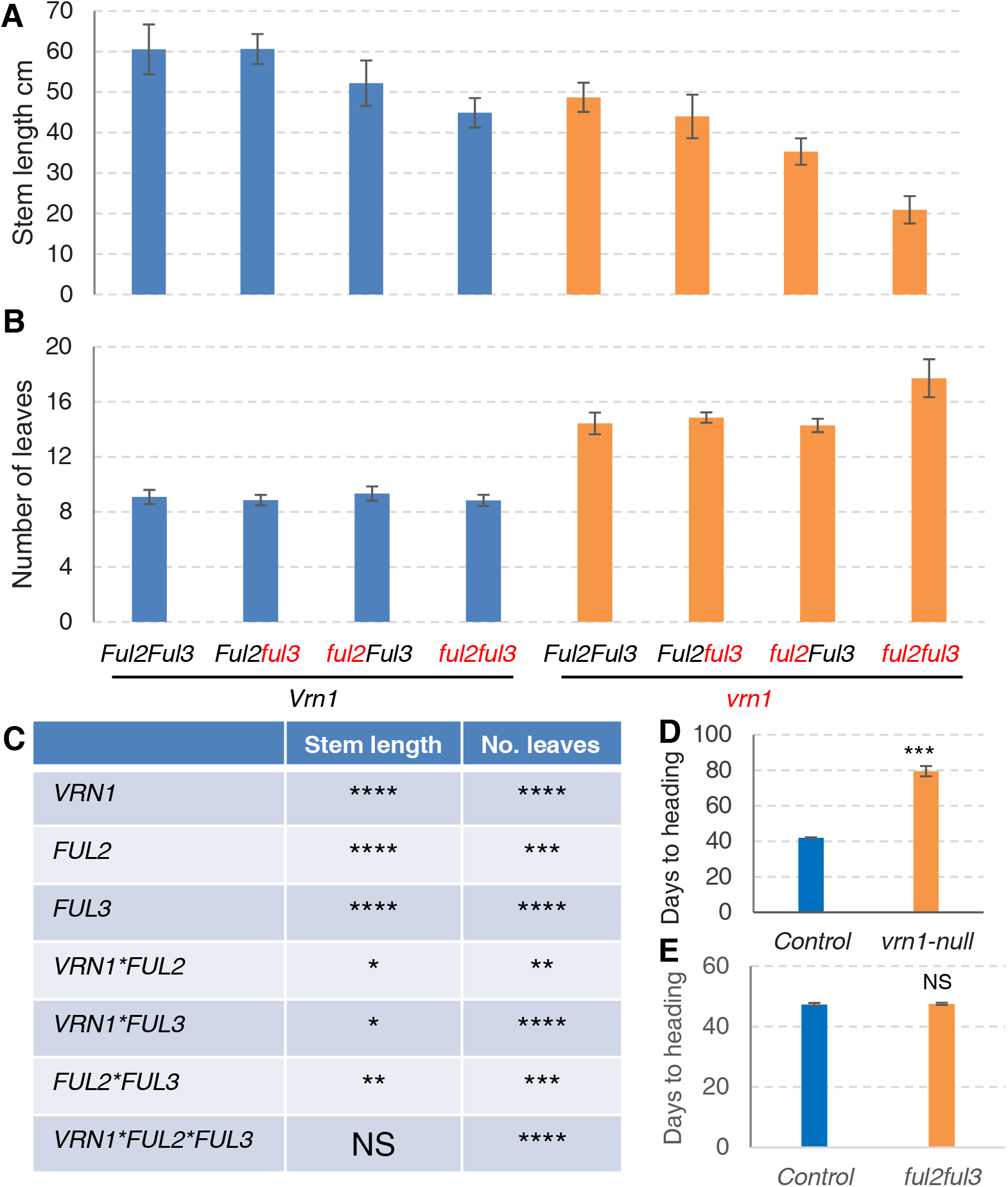
Effect of *VRN1, FUL2* and *FUL3* on stem length, leaf number and heading time. Kronos plants (*vrn2*-null background) grown under long-day photoperiod. Stem length was determined from the base of the plant to the base of the spike. (A) Stem length in cm (n= 6-12). (B) Number of true leaves (n= 6-12). Alleles in red indicate homozygous null mutants and alleles in black homozygous wild type alleles. (C) *P* values from three-way ANOVAs for stem length and leaf number including all eight homozygous *VRN1*, *FUL2* and *FUL3* allele combinations (n= 59). * = *P* < 0.05, ** = *P* < 0.01, *** = *P* < 0.001, **** = *P* < 0.0001, NS = *P* > 0.05. (D) Heading time of *vrn1*-null (n=6) versus control (n= 6). (E) Heading time of *ful2ful3*-null (n= 15) *vs*. control (n= 10) in a *Vrn1* background. D and E are separate experiments. Error bars are SEM.

### *VRN1*, *FUL2* and *FUL3* loss-of-function mutations reduce stem elongation

Since some mutant combinations lack real spikes, we determined final stem length from the base of the plant to the base of the spike (or spike-like structure) instead of total plant height. Plants carrying only the *ful3*-null mutation showed no significant reduction in stem length, but those carrying the *vrn1*-null or *ful2*-null mutations were 20% and 14% shorter than the control, respectively (Fig. 1A). A three-way factorial ANOVA for stem length revealed highly significant effects for all three genes (Fig. 1C). All three double mutant combinations had shorter stems than predicted from combined additive effects of the individual mutations, which was reflected in significant synergistic interactions (Fig. 1C). Taken together, these results indicate that *VRN1*, *FUL2* and *FUL3* have redundant roles in the regulation of stem elongation, and that the effect of the individual genes is larger in the absence of the other paralogs.

### *VRN1*, *FUL2* and *FUL3* mutations delay flowering time

Functional redundancy among *VRN1*, *FUL2* and *FUL3* was also observed for heading time. The *vrn1*-null mutant headed 37.5 d later than the control (Fig. 1D), but differences in heading time for the *ful2*-null, *ful3*-null and *ful2ful3*-null mutants in the presence of the strong *Vrn-A1* allele were non-significant (Fig. 1E). For the *vrn1ful2*-null and *vrn1ful2ful3*-null mutants, it was not possible to determine heading times accurately because they had short stems and abnormal spikes that interfere with normal ear emergence. Instead, we determined the final number of leaves (Fig. 1B) and the timing of the transition between the vegetative and double-ridge stages (Fig. S3).

A three-way factorial ANOVA for leaf number revealed highly significant effects for the three individual genes, as well as for all the two- and three-way interactions (Fig. 1C). The *vrn1*-null mutant had on average 14.4 leaves (59% > control, Fig. 1B), which was consistent with its later heading time (Fig. 1D). Similar leaf numbers were detected in *vrn1ful2*-null (14.3) and *vrn1ful3*-null (14.9), but the triple *vrn1ful2ful3*-null mutant had on average 17.7 leaves (Fig. 1B), which was consistent with the 9 to 12 d delay in the transition between the vegetative SAM and the double-ridge stage relative to the *vrn1-null* control (Fig. S3). These results indicate that *FUL3* has a residual ability to accelerate flowering time in the absence of *VRN1* and *FUL2*.

Transgenic Kronos plants overexpressing the coding regions of *FUL2* fused with a C-terminal 3xHA tag (henceforth *Ubi::FUL2*, Fig. S4A, events #1 and #6) or *FUL3* fused with a C-terminal 4xMYC tag (henceforth *Ubi::FUL3*, Fig. S4B, events #4 and #5) headed two to four days earlier than the non-transgenic sister lines (*P* < 0.0001). The effect of *Ubi::FUL2* was further characterized in the F_2_ progeny from the cross between *Ubi::FUL2* (*Vrn1Vrn2*) and *vrn1vrn2*-null under greenhouse conditions. A three-way factorial ANOVA for heading time showed significant effects for *VRN1*, *Ubi:FUL2* and *VRN2* and for all two- and three-way interactions (*P* < 0.0001, Table S3). In the presence of a functional *VRN2* allele, the differences in heading time between *FUL2-wt* and *Ubi::FUL2* alleles were small in lines homozygous for the functional *VRN1* allele (2.6 d, Fig. S4A), intermediate in *VRN1* heterozygous lines (11.1 d, Fig. S4C) and large in homozygous *vrn1-null* mutants (53 d, Fig. S4D). These results indicate that the effect of the *Ubi::FUL2* transgene on heading time depends on the particular *VRN1* and *VRN2* alleles present in the genetic background (Fig. S4C-D).

In summary, the strong effect of *VRN1* in the acceleration of wheat flowering time can mask the smaller effects of *FUL2* and *FUL3*, but in the absence of *VRN1*, both *FUL2* and *FUL3* have redundant effects on accelerating wheat flowering time.

### Flowering delays in *VRN1*, *FUL2* and *FUL3* mutants are associated with reduced *FT1* transcript levels in leaves

Since there is a known positive feedback regulatory loop between *VRN1* and *FT1* (Shaw et al., 2019), we compared the *FT1* transcript levels in the leaves of the different *VRN1*, *FUL2* and *FUL3* mutant combinations. *FT1* transcript levels higher than *ACTIN* were observed in leaves of 4-week old plants carrying the *VRN1* wild type allele, but were detected only after 10 weeks in plants carrying the *vrn1*-null allele (Fig. S5A-B). This result is consistent with the large differences in heading time between these genotypes (Fig. 1D). *FT1* transcript levels in the 10-week old *vrn1*-null plants was highest in the presence of the *FUL2* and *FUL3* wild type alleles and lowest in the triple mutant (Fig. S5C), consistent with the higher number of leaves in this genotype (Fig. 1B). Even in the *vrn1ful2ful3*-null plants, *FT1* transcript levels increased above *ACTIN* in 14-week old plants (Fig. S5D). Taken together, these results indicate that *FT1* expression in the leaves are positively regulated by *VRN1*, *FUL2* and *FUL3*, but that they can also be up-regulated in the absence of these three genes.

### *VRN1, FUL2,* and *FUL3* play critical and redundant roles in spikelet development

Plants with individual *vrn1*-null, *ful2*-null and *ful3*-null mutations produced normal spikelets and flowers, but *vrn1ful2*-null or *vrn1ful2ful3*-null mutants had spike-like structures in which all lateral spikelets were replaced by leafy shoots (Fig. 2A-J), henceforth referred to as “inflorescence tillers”. Removal of these inflorescence tillers revealed a thicker and shorter rachis with fewer internodes of variable length, but still retaining the characteristic alternating internode angles typical of a wild type rachis (Fig. 2B).

**Fig. 2.**
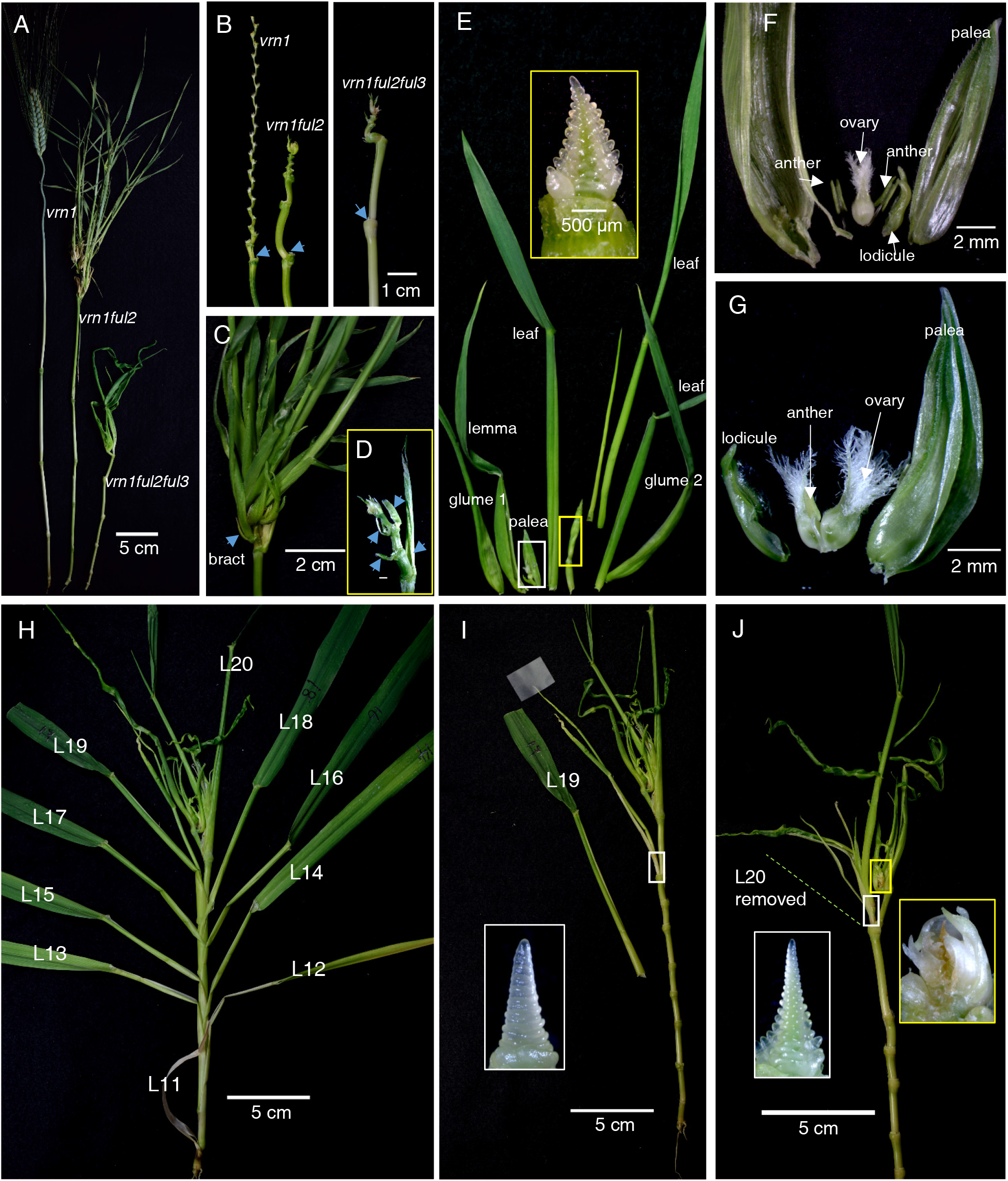
Phenotypical characterization of the *vrn1ful2* and *vrn1ful2ful3* mutants. (A) Stems and heads of *vrn1*-null, *vrn1ful2*-null and *vrn1ful2ful3*-null mutants (leaves were removed before photography). (B) Rachis of the different mutants. Arrows indicate the position of the first spikelet before removal. (C-G) *vrn1ful2*-null mutant. (C) Spike-like structure. Arrow points to the bract subtending a basal inflorescence tiller. (D) Spike-like structure after removal of the inflorescence tillers to show subtending bracts (arrows). (E) Dissection of an inflorescence tiller showing two glumes and one lemma partially transformed into leaves, followed by four leaves. The inset with yellow border shows the meristem transition into an IM with lateral VMs. (F) Detail of white rectangle in (E) revealing an ovary, two anthers, and leafy-lemma and palea. (G) Leafy palea and lodicules subtending one anther and two ovaries. (H-J) *vrn1ful2ful3*-null mutant. (H) Normal leaves L11 to L18 with no axillary buds. L19 marks the beginning of the spike-like structure in which spikelets have been replaced by tillers subtended by leaves (L19 and L20) or bracts. (I and J) Detail of the tillers subtended by L19 (I) and L20 (J). Insets in white rectangles are the SAM of these tillers (transitioning into IM with lateral VM) and the yellow rectangle presents the exhausted IM.

In *vrn1ful2*-null, approximately 70% of the central inflorescence tillers had leafy glumes, lemmas and paleas and abnormal floral organs, whereas the rest were fully vegetative. Floral abnormalities included leafy lodicules, reduced number of anthers, anthers fused to ovaries, and multiple ovaries (Fig. 2E-G). After the first modified floret, meristems from these inflorescence tillers developed two to five true leaves before transitioning again to an IM generating lateral VMs (Fig. 2E). The combined presence of floral organs and leaves suggests that the originating meristem had an intermediate identity between VM and SM. In the *vrn1ful2*-null double mutant the inflorescence tillers were subtended by bracts (Fig. 2C-D).

In *vrn1ful2ful3*-null, the lateral meristems generated inflorescence tillers that had no floral organs, and that were subtended by leaves in the basal positions and bracts in more distal positions (Fig. 2H-J). The presence of well-developed axillary tillers in these basal inflorescence leaves (Fig. 2H, L19 and L20) marked the border of the spike-like structure, because no axillary tillers or developing buds were detected in the true leaves located below this border (Fig. 2H, L11-L18).

Scanning Electron-Microscopy (SEM) images of the early developing inflorescences in the *vrn1ful2*-null and *vrn1ful2ful3*-null mutants revealed elongated double-ridge structures similar to those in Kronos (Fig. 3 A) or *vrn1*-null (Fig. 3 C). Suppression of the lower ridge was complete in Kronos (Fig. 3A) and in *vrn1*-null (Fig. 3D, red arrows), but was incomplete in *vrn1ful2*-null (Fig. 3B, E; yellow arrows), and even weaker in *vrn1ful2ful3*-null (Fig. 3C, F: green arrows). As a result of this change, inflorescence tillers were subtended by bracts in *vrn1ful2*-null (Fig. 2C-D) and by leaves in *vrn1ful2ful3*-null (Fig. 2H-I). The upper ridges (Fig. 3A-C, dots) transitioned into normal SMs in *vrn1*-null, with glume and lemma primordia (Fig. 3D, G), but looked like typical vegetative meristems in *vrn1ful2*-null and *vrn1ful2ful3*-null (Fig. 3E-F, H-I).

**Fig. 3.**
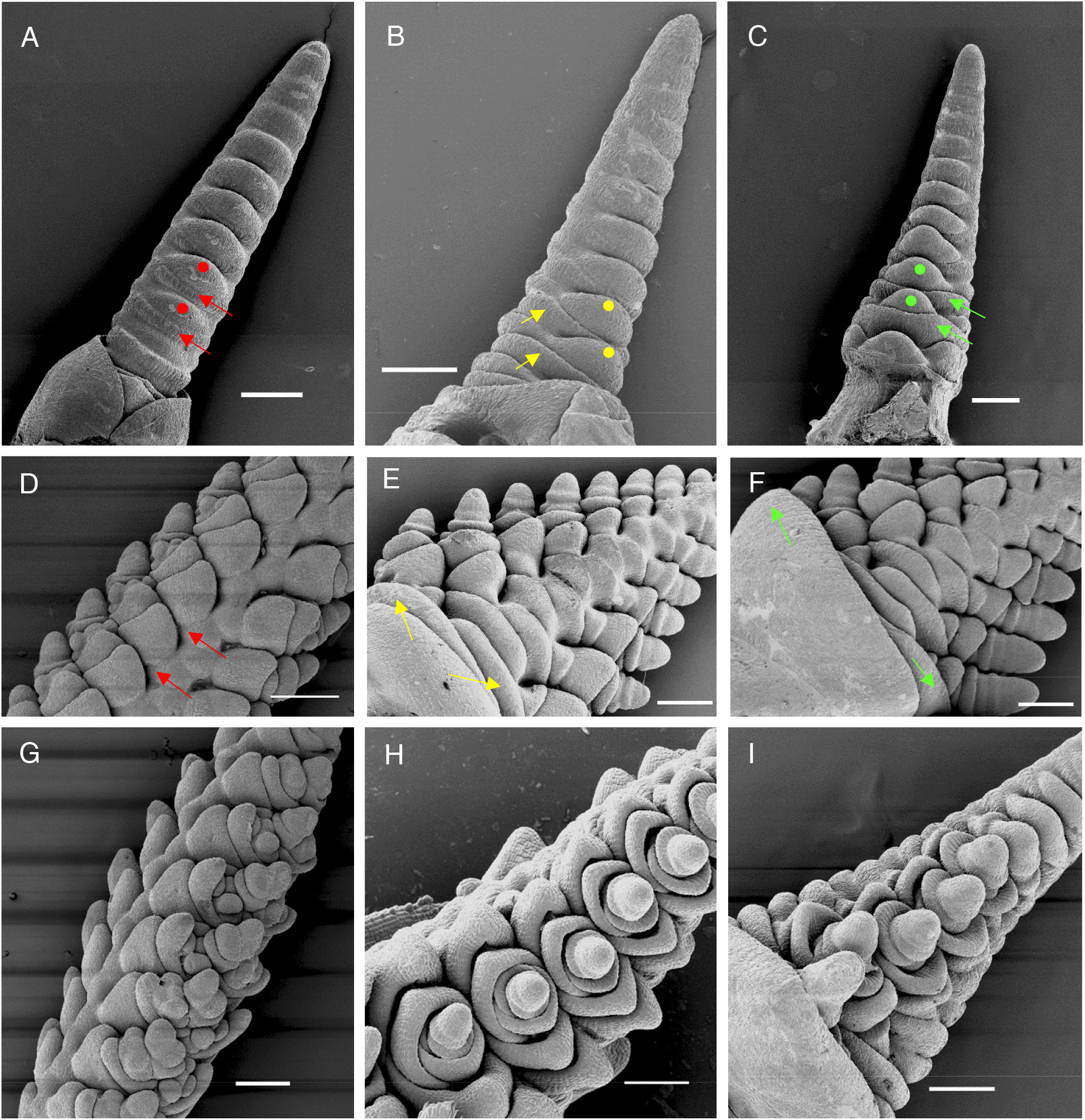
Scanning Electron Microscopy (SEM) images. (A-C) Early double-ridge stage (D-I) Later stage showing the fate of the lateral meristems. (A) Kronos control (D, G) *vrn1*-null control. Red arrows indicated the repressed lower ridge, and red dots the upper ridges that develop into normal spikelets (D and G). (B, E, H) *vrn1ful2*-null mutants. Yellow arrows indicate the partially repressed lower ridges that develop into bracts (see Fig. 2D) and yellow dots indicate the upper ridges that develop into intermediate meristems that generate tiller-like structures with altered floral organs (see Fig. 2E-G). (C, F, I) *vrn1ful2ful3*-null mutants. Green arrows indicate basal lower ridges that develop into normal leaves (see Fig. 2H) and green dots indicate upper ridges that produce lateral vegetative meristems that generate vegetative tillers with no floral organs (see Fig. 2 I-J). Bar = 200 μm.

To characterize better the relative effects of *VRN1* and *FUL2*, we examined their individual effects when present as a single functional copy in a heterozygous state (underlined). Both *ful2-null/<u>Vrn-A1</u> vrn-B1-null* (functional *Vrn1* allele for spring growth habit) and *ful2-null/vrn-A1-null <u>vrn-B1</u>* (functional *vrn1* allele for winter growth habit) produced spike-like structures with leafy lateral shoots (Fig. S6A-B) and normal floral organs (Fig. S6C) but no viable seeds. The developing spikes of these plants showed lateral meristems with floral primordia (Fig. S6D), some of which later showed elongated rachillas and leafy organs (Fig. S6E-G). By contrast, the presence of a single heterozygous copy of *FUL2 (vrn1-null/ful2-A <u>Ful2-B</u>*) was sufficient to generate more normal-looking spikelets (Fig. S6H-J), some of which were able to set viable seeds. Abnormal spikelets (Fig. S6I) and basal branches with lateral spikelets and fertile florets (Fig. S6J) were also observed in this mutant. Taken together these results indicate that *VRN1, FUL2* and *FUL3* have redundant and essential roles in spikelet development, with *FUL2* having the strongest effect and *FUL3* the weakest.

### Transcript levels of *SHORT VEGETATIVE PHASE* (*SVP*)-like genes *VRT2, BM1* and *BM10* are upregulated in the developing spikes of the *vrn1ful2*-null mutant

A partial reversion of basal spikelets to vegetative tillers similar to the one described above for the *vrn1ful2*-null mutant, has been described in barley lines overexpressing *SVP*-like genes *BM1* or *BM10* (see Discussion). To test if the transcript levels of the *SVP*-like wheat genes were affected in the *vrn1ful2*-null mutants, we first determined their expression during normal spike development in Kronos. Transcript levels of the three related paralogs *BM1, BM10* and *VRT2* (RefSeq v1.0 gene designations in Fig. S7) decreased three-to five-fold from the initial stages of spike development (W2, Waddington scale) to the floret primordium stage (W3.5, Fig. S7A-C).

Next, we compared the transcriptional levels of the *SVP*-like wheat genes in *vrn1ful2*-null and *vrn1*-null mutants. Plants were grown for 53 days in a growth chamber until the developing spikes of *vrn1*-null reached the terminal spikelet stage and those from *vrn1ful2*-null had a similar number of lateral meristems. The transcript levels of *BM1, BM10* and *VRT2* in the developing spikes were roughly ten-fold higher in the *vrn1ful2*-null mutant than in the *vrn1*-null and control lines (*P* < 0.0001, Fig. S7D-F). These results suggest that *VRN1* and *FUL2* are either direct or indirect transcriptional repressors of the three wheat *SVP*-like genes.

### *FUL2* and *VRN1* have redundant roles on spike determinacy and regulate the number of spikelets per spike

Normal wheat spikes are determinate, with the distal IM transitioning into a terminal spikelet after producing a relatively stable number of lateral meristems (Fig. 4A). In *vrn1ful2*-null, by contrast, the IM was indeterminate (Fig. 4B) and continued to produce lateral meristems while growing conditions were favorable and eventually died without producing any terminal structure. In the *ful2*-null background, one functional copy of *VRN1* in the heterozygous state was sufficient to generate a determinate spike (Fig. S6D, *ful2*-null/*vrn-A1*-null <u>*vrn-B1*</u>), and the same was true for a single functional copy of *FUL2* in a *vrn1*-null background (Fig. S6K, *vrn1*-null/*ful2-A <u>Ful2-B</u>*).

**Fig. 4.**
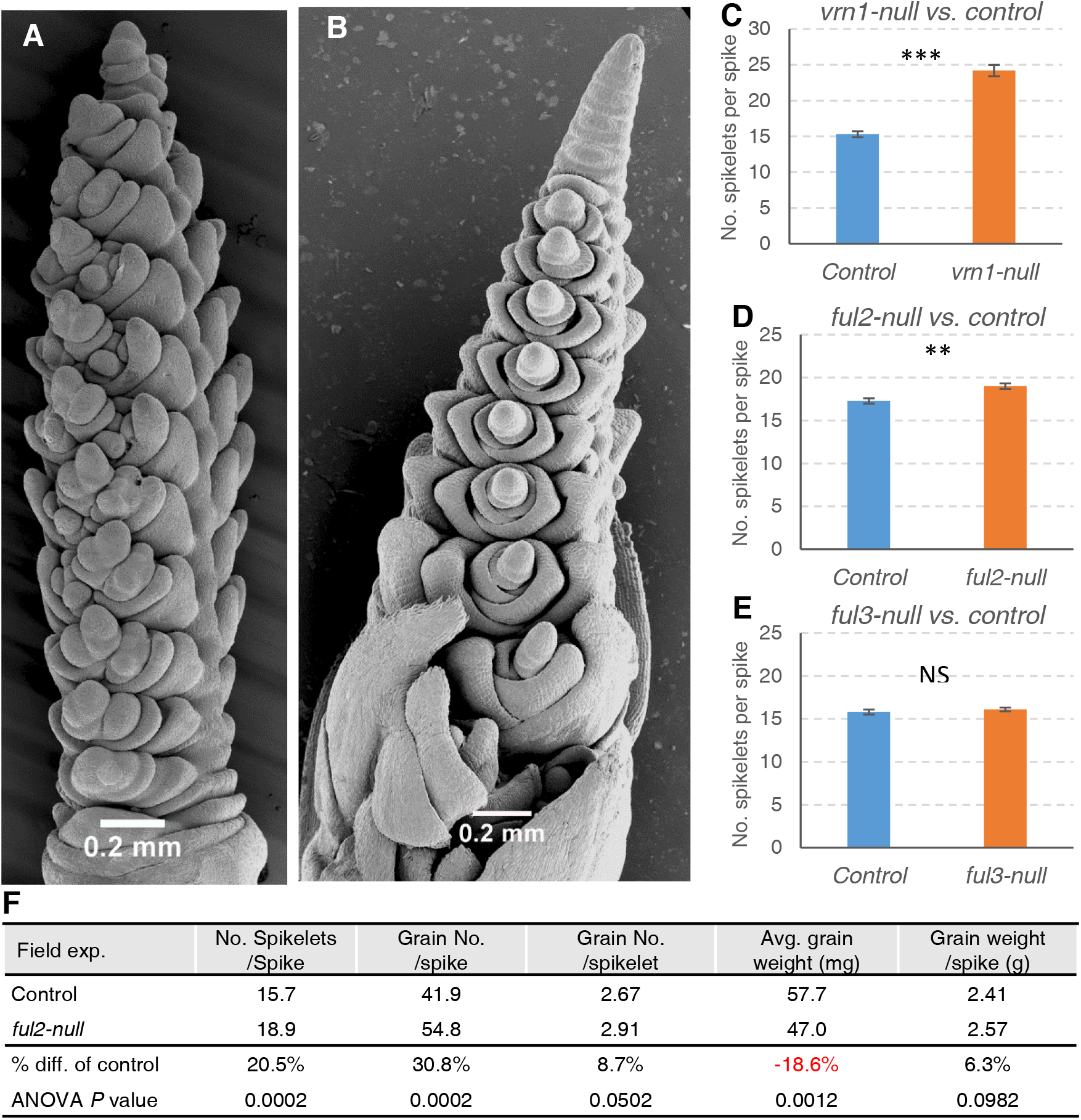
*VRN1* and *FUL2* play redundant roles in the control of spike determinacy and spikelet number. (A) Scanning Electro-Microscopy of a normal wheat spike with a terminal spikelet in *vrn1*-null. (B) *vrn1ful2*-null mutant spike with indeterminate apical meristem. (C-E) Number of spikelets per spike in a growth chamber experiment (n=6). (C) *vrn1*-null (58% increase), (D) *ful2*-null (10% increase) and (E) *ful3*-null (no significant increase). Bars represent mean ± SEM and asterisks indicate statistically significant difference to the control line (** = *P* < 0.01, *** = *P* <0.001, NS = *P* > 0.05). (F) ANOVAs for spike traits in *ful2*-null and sister control lines in the field (randomized complete block design with 8 blocks).

The individual *vrn1*-null and *ful2*-null homozygous mutants showed a larger number of spikelets per spike than the control. This increase was 58% in the *vrn1*-null mutant (*P* < 0.0001, Fig. 4C) and 10% in the *ful2*-null mutant (*P* = 0.0014, Fig 4D). Although no significant increases in the number of spikelets per spike were detected in the *ful3*-null mutant (*P* = 0.4096, Fig. 4E), two independent transgenic lines overexpressing *FUL3* (*Ubi::FUL3*) showed an average reduction of 1.12 spikelet per spike relative to their non-transgenic sister lines (*P* = 0.0132 and *P* < 0.0001, Fig. S8A), which indicates that *FUL3* can still play a role on the timing of the transition from IM to terminal spikelet.

A similar reduction in the number of spikelets per spike was observed in two independent *Ubi::FUL2* transgenic lines (1.05 spikelets per spike reduction, *P* <0.03, Fig. S8B). We then investigated the effect of this transgene in the presence of different *VRN1* and *VRN2* alleles in the *Ubi::FUL2* x *vrn1vrn2*-null F_2_ population. In the *vrn2*-null F_2_ plants, the differences in spikelet number between *Ubi::FUL2* and wild type alleles were larger in *vrn1*-null than in the *Vrn1*-Het plants (interaction *P* < 0.0001, Fig. S8C). In a separate group of F_2_ plants fixed for *Vrn1*-Het and segregating for *VRN2* and *FUL2*, we did not detect significant effects for *Ubi::FUL2* and the interaction was not significant (Fig. S8D). However, we observed 3.3 more spikelets per spike in *Vrn2*-wt than in *vrn2*-null plants (*P* < 0.0001, Fig. S8D). These results suggest that the strong *Vrn-A1* allele for spring growth habit can mask the effects of the *Ubi::FUL2* transgene but not that of *VRN2* on the number of spikelets per spike.

### Increased transcript levels of *CEN2, CEN4* and *CEN5* in developing spikes of the *vrn1ful2*-null mutant

Based on the strong effect observed in the Arabidopsis *tfl1* mutant and the *Antirrhinum cen* mutant on inflorescence determinacy (see Discussion), we investigated the effect of the *vrn1ful2*-null mutations on the expression levels of the *TFL1/CEN*-like wheat homologs in developing spikes. Since no previous nomenclature was available for the wheat *CEN* paralogs, we assigned them numbers to match their chromosome locations, and designated them as *CEN2, CEN4* and *CEN5* (RefSeq v1.0 designations can be found in the legend of Fig. S9). The transcript levels of these three genes were downregulated as the developing spike progressed from the double-ridge stage to the floret primordium stage (Waddington scale 2 to 3.5, Fig. S9A-C).

Comparison of *vrn1ful2*-null and *vrn1*-null plants grown for 53 days in the same growth chamber showed that the transcript levels of *CEN2, CEN4* and *CEN5* were significantly higher *(P* < 0.0001) in the developing spikes of the *vrn1ful2*-null mutant than in those of the *vrn1*-null mutant or the Kronos control (all in *vrn2*-null background). These differences were larger for *CEN2* and *CEN4* than for *CEN5* (Fig. S9D-F). Taken together, these results suggested that *VRN1* and *FUL2* work as transcriptional repressors of the *TFL1/CEN*-like wheat homologs.

### The *ful2*-null mutant produces a higher number of florets per spikelet and more grains per spike in the field

In addition to the higher number of spikelets per spike, the *ful2*-null mutant produced a higher number of florets per spikelet than the Kronos control, an effect that was not observed for *vrn1*-null (Fig. 2A) or *ful3*-null (Fig. S10A). The average increase in floret number was similar in *ful2*-null (1.3 florets) and *ful2ful3*-null (0.9 florets), suggesting that *FUL3* has a limited effect on this trait. In spite of some heterogeneity in the distribution of spikelets with extra florets among spikes, the differences between the control and the *ful2*-null mutants were significant at all spike positions (Fig. S10B).

Similar increases in the number of florets per spikelet were reported before in Kronos plants overexpressing *miRNA172* under the *UBIQUITIN* promoter (Debernardi et al., 2017). To study the genetic interactions between *Ubi::miR172* and *ful2*-null we crossed the transgenic and mutant lines and studied their effects on floret number in the progeny using a two-way factorial ANOVA. We detected significant differences in average floret number for both *ful2*-null and *Ubi::miR172* (*P* < 0.01) and a marginally significant interaction (*P* < 0.0435) that can be visualized in the interaction graph in Fig. S10C. The differences in average floret number between *ful2*-null and the wild type control were larger (and more variable) in the *Ubi::miR172* than in the non-transgenic background (Fig. S10C). This synergistic interaction suggests that *miRNA172* and *FUL2* may control floret number through a common pathway. Both the mutant and transgenic lines showed heterogeneity among spikes in the location of spikelets with increased numbers of florets (Fig. S10D-F).

Based on its positive effect on the number of florets per spikelet and spikelets per spike (and its small effect on heading time), we selected the *ful2*-null mutant for evaluation in a replicated field experiment. Relative to the control, the *ful2*-null mutant produced 20% more spikelets per spike (*P* = 0.0002) and 9% more grains per spikelet (*P*= 0.05), which resulted in a 31% increase in the number of grains per spike (*P* = 0.0002, Fig. 4F). Although part of the positive effect on grain yield was offset by a 19% reduction in average kernel weight (*P* = 0.0012), we observed a slight net increase of 6% in total grain weight per spike (*P* = 0.09, Fig. 4F). This negative correlation between grain number and grain weight suggests that in this particular genotype by environment combination grain yield was more limited by the “source” (produced and transported starch) than by the “sink” (number and size of grains).

## DISCUSSION

Results from this study have shown that wheat *VRN1*, *FUL2* and *FUL3* have overlapping functions in stem elongation, flowering time, and spike development, which are discussed separately in the following sections.

### Mutations in *VRN1*, *FUL2* and *FUL3* reduce stem elongation

We detected highly significant effects of *VRN1*, *FUL2* and *FUL3* on plant height and significant synergistic interactions (Fig. 1A, C). These results suggest that *VRN1*, *FUL2* and *FUL3* have redundant functions in the regulation of stem elongation, and that their individual effects are magnified in the absence of the other paralogs. Significant reductions in plant height were also reported for rice mutants *Osmads14* (12.2% reduction) and *Osmads15* (9.0 % reduction), and the double mutant (43.8% reduction), suggesting a conserved function in grasses (Wu et al., 2017).

Although the molecular mechanisms by which these genes affect stem elongation are currently unknown, an indirect way by which they may contribute to this trait is through their strong effect on the regulation of *FT1* (Fig. S5), which is associated with the upregulation of GA biosynthetic genes in the developing spike (Pearce et al., 2013). A recent study has shown that rice *HEADING DATE 3* (*Hd3*) and *RICE FLOWERING LOCUS T 1* (*RFT1*), the orthologs of wheat *FT1*, can increase stem responsiveness to GA by reducing *PREMATURE INTERNODE ELONGATION 1* (*PINE1*) expression in the SAM and compressed stem (Gómez-Ariza et al., 2019). In Arabidopsis, a number of genes involved in hormone pathways are direct targets of FUL, which may explain the shorter stem and internodes detected in the *ful* mutant (Bemer et al., 2017). A characterization of the direct DNA targets and protein interactors of VRN1, FUL2 and FUL3 may shed light on the mechanisms responsible for the conserved role of these genes in plant height in grasses.

### Mutations in *VRN1*, *FUL2* and *FUL3* delay flowering initiation in wheat

*VRN1* is one of the main genes controlling natural variation in wheat flowering time (Fu et al., 2005; Kippes et al., 2016; Yan et al., 2003; Zhang et al., 2008), so it was not surprising that *vrn1*-null delayed heading time more than *ful2*-null or *ful3*-null. Although the strong *Vrn-A1* allele for spring growth habit masked the smaller effects of *FUL2* and *FUL3* (Fig. 1A-C), in the *vrn1*-null background, the *ful2*-null and *ful3*-null mutants showed delayed flowering initiation and increased number of leaves (Fig. 1B, F), indicating that *FUL2* and *FUL3* have retained some residual functionality in the acceleration of wheat flowering time. This was further confirmed by the accelerated flowering of the *Ubi::FUL2* and *Ubi::FUL3* transgenic plants (Fig. S4A-B). Similar results have been reported in *Brachypodium distachyon* and rice. In *Brachypodium*, overexpression of *VRN1* (Ream et al., 2014), *FUL2* or *FUL3* (Li et al., 2016) accelerates flowering, and downregulation of *VRN1* delays flowering relative to non-transgenic controls (Woods et al., 2016). In rice, overexpression of *OsMADS15* also accelerates flowering (Lu et al., 2012). These results suggest a conserved role of these genes in the regulation of flowering time in grasses.

Previous studies have shown a significant genetic interaction between wheat *VRN1* and *VRN2* in the regulation of heading time (Tranquilli and Dubcovsky, 2000). This study shows that similar interactions exist between *FUL2* and *VRN2* (Fig. S4C-D). A tetraploid wheat population segregating for *VRN1*, *FUL2* and *VRN2* revealed highly significant two-way and three-way interactions among these genes, indicating that the effect of each of these genes on heading time is dependent on the particular combination of alleles present for the other two. Previous studies have shown that part of the ability of *VRN1* to accelerate flowering depends on its ability to repress *VRN2* (Chen and Dubcovsky, 2012). The larger effect on heading time of the *Ubi::FUL2* transgene in the presence of the functional *Vrn2* than in the *vrn2*-null background (Fig. S4C-D) suggests that *FUL2* repression of *VRN2* can also contribute to its ability to accelerate heading time.

Interestingly, mutations in *VRN1*, *FUL2* and *FUL3* were associated with delayed induction of *FT1* even in the absence of functional *VRN2* alleles (Fig. S5). Lines carrying the wild type *VRN1* allele showed high levels of *FT1* in the leaves six weeks earlier than lines carrying the *vrn1*-null allele and flowered 37 days earlier. Among the lines carrying the *vrn1*-null allele, those with mutation in both *FUL2* and *FUL3*, showed the latest induction of *FT1* (Fig. S5C-D) and had 3.3 more leaves (excluding leaves in the spike-like structure, Fig. 1B). However, in 14-week-old *vrn1ful2ful3*-null plants *FT1* transcripts still reached higher levels than *ACTIN*. These results indicate that VRN1, FUL2 and FUL3 are positive transcriptional regulators of *FT1* but they are not essential for its expression in the leaves.

FT1 has also been shown to be a positive regulator of *VRN1* expression in both leaves and SAM. Natural variation or transformation experiments that affect *FT1* transcript levels in the leaves are always associated with parallel changes in *VRN1* expression (Lv et al., 2014; Yan et al., 2006). Taken together, these results suggest the existence of a positive regulatory feedback loop in which each gene acts as a positive regulator of the other. Although this feedback loop can be mediated in some cases by *VRN2* (Distelfeld et al., 2009a), results from this study and from Shaw et al. (2019) in a *vrn2*-null background suggest the existence of a positive feedback loop that operates independently of *VRN2*. Studies in rice have suggested the possibility of a similar regulatory feedback loop between the orthologous genes (Kobayashi et al., 2012).

### *VRN1*, *FUL2* and *FUL3* play critical and redundant roles in spikelet development

RNA *in situ* hybridization studies at the early stages of inflorescence development in *Lolium temulentum* (Gocal et al., 2001), wheat and oat (Preston and Kellogg, 2008), and early-diverging grasses (Preston et al., 2009) have shown *VRN1* and *FUL2* expression in the IM, lateral SMs and FMs. Similarly, transgenic barley plants transformed with a *VRN-H1* promoter fused with GFP showed fluorescence in the three meristems (Alonso-Peral et al., 2011). VRN1, FUL2, and FUL3 can interact with different MADS-box partners in the IM, SMs, and FMs and, therefore, mutations in these genes can alter different functions in different meristems.

The significant effects of *VRN1*, *FUL2*, and *FUL3* on the timing of the transitions from the vegetative SAM to IM and from IM to terminal spikelet indicate that these genes play important roles in the acquisition and termination of IM identity. During the early stages of spike development both *vrn1ful2*-null and *vrn1ful2ful3*-null mutants showed an elongated double-ridge structure with lateral meristems organized in a distichous phyllotaxis that were similar to the Kronos control (Fig. 3A-C), and both mutants had a rachis similar to a normal spike rachis (Fig. 2B). These results suggest that these IM functions were not disrupted by the combined mutation in *VRN1*, *FUL2*, and *FUL3*.

After the double-ridge stage, the development of the lateral meristems diverged drastically in *vrn1ful2*-null and *vrn1ful2ful3*-null mutants relative to the *vrn1*-null control. In *vrn1*-null, the upper ridges transitioned into SMs that generated normal spikelets, whereas in *vrn1ful2ful3*-null they transitioned into lateral VMs that generated inflorescence tillers (which we interpret as the default identity of an axillary meristem). The *vrn1ful2*-null mutant generated an intermediate structure that produced both leafy-floral organs and leaves. Based on these results we concluded that *VRN1, FUL2* and *FUL3* play essential and redundant roles in spikelet and floral development. However, we currently do not know if the transition of the upper ridges to SMs requires the expression of functional VRN1, FUL2 and FUL3 proteins in the upper ridge, in the IM, or in both.

Replacement of basal spikelets with inflorescence tillers similar to the ones described for *vrn1ful2ful3*-null was observed in barley plants overexpressing *BM1* and *BM10* (Trevaskis et al., 2007). These two genes, together with *VRT2*, are related to the Arabidopsis MADS-box genes *SVP* and *AGAMOUS-LIKE 24* (*AGL24*), which play important roles in the formation of floral meristems (Kaufmann et al., 2010; Liu et al., 2007). In Arabidopsis, *SVP* and *AGL24* are directly repressed by AP1 (Kaufmann et al., 2010). In the absence of AP1, ectopic expression of *SVP* and *AGL24* transformed floral meristems into shoot meristems (Liu et al., 2007). The up-regulation of *VRT2*, *BM1* and *BM10* in the *vrn1ful2*-null developing wheat spikes (Fig. S7), together with the transgenic barley results, suggest that these genes may have contributed to the observed replacement of spikelets by vegetative tillers in *vrn1ful2*-null.

### Both *vrn1ful2*-null and *vrn1ful2ful3*-null mutants showed reduced suppression of the lower ridge

An important characteristic of a grass IM is the complete suppression of the lower ridge subtending all branching events, which in the wheat spike is the suppression of the lower ridge subtending the spikelet. This suppression was disrupted in the *vrn1ful2*-null and *vrn1ful2ful3*-null mutants, which developed bracts or leaves subtending the inflorescence tillers (Fig. 2C-D and H-J). These results suggest that all three genes contribute to the suppression of the lower ridge, but we do not know if this suppression requires the expression of the VRN1, FUL2 and FUL3 in the lateral meristem, in the IM, or in both. In this case, an indirect IM effect seems more likely because *in situ* hybridization studies have detected *VRN1* and *FUL2* (or their grass orthologs) in the upper ridge rather than in the lower ridge (Gocal et al., 2001; Preston and Kellogg, 2008) (Preston et al., 2009). However a more direct effect on the lateral meristem cannot be ruled out because a VRN1:GFP fusion driven by the *VRN1* promoter was detected in the lower ridge of the developing spike in barley (Alonso-Peral et al., 2011).

The inflorescence tillers subtended by leaves in *vrn1ful2ful3*-null were not very different from vegetative tillers, but there was a difference that marked a clear boundary between them. In rice and wheat, true leaves do not show axillary buds or tillers until 4-5 younger leaves are formed (Friend, 1965; Oikawa and Kyozuka, 2009). Then, bud development into tillers proceeds sequentially from older to younger leaves. By contrast, leaves developed from the lower ridge in *vrn1ful2ful3*-null (Fig. 3C, F) had axillary meristems (the upper ridge) from the beginning, which rapidly developed into axillary tillers (Fig. 2H, L19 and L20). True leaves below the inflorescence (Fig. 2H, L11-L18) showed no visible axillary buds, which is normal for wheat leaves that subtend an elongated internode (Williams and Langer, 1975). In summary, even in the *vrn1ful2ful3*-null mutant a clear boundary was established between the inflorescence and the vegetative leaves.

### Interpretation of observed meristem identity changes

An inflorescence phenotype similar to the one described here for the *vrn1ful2ful3*-null wheat mutant has been described for the rice *Osmads14Osmads15Osmads18Ospap2* quadruple knock-down, in which the panicle was replaced by tillers subtended by leaves (Kobayashi et al., 2012). The authors of the rice study interpreted this phenotype as the result of an incomplete transition between the vegetative SAM and the IM, and suggested that these genes act redundantly to promote IM identity and, therefore, are IM identity genes.

In wheat *vrn1ful2ful3*-null, the changes observed in the lateral meristems can be also explained by postulating an indirect effect of the IM on the regulation of genes expressed in the signaling centers flanking the lateral meristems. However, the same changes can be explained by a more direct effect of non-functional VRN1, FUL2 and FUL3 proteins in the lateral meristem, where they are normally expressed. If this second interpretation is correct, *VRN1*, *FUL2* and *FUL3* should be considered to include SM identity functions, in addition to IM and FM identity functions. This proposed SM identity function is consistent with the role of homologous FM identity genes *AP1*, *CAL* and *FUL* in Arabidopsis (Ferrándiz et al., 2000). Regardless of their direct or indirect effect on SM identity, *VRN1*, *FUL2* and *FUL3* are essential for spikelet development in both wheat and rice.

This does not seem to be the case for *PAP2* or its wheat ortholog *AGLG1* (Yan et al., 2003). Kobayashi et al. (2012) suggested that the loss of function of this gene was important for the rice *Osmads14Osmads15Osmads18Ospap2* phenotype. However, the complete suppression of spikelets in the presence of functional *PAP2/AGLG1* genes in rice *Osmads14Osmads15* (Wu et al., 2017) and wheat *vrn1ful2ful3*-null mutants suggests a less critical role of *PAP2* on SM identity.

### *VRN1* and *FUL2* have essential and redundant roles in wheat spike determinacy

The determinate growth of the wheat spike is marked by the transition of the distal IM into a SM and the formation of a terminal spikelet. However, the *vrn1ful2*-null mutant was unable to form spikelets and the IM remained indeterminate. A single functional copy of *VRN1* or *FUL2* in a heterozygous state was sufficient to restore spike determinacy (Fig S5D, K), suggesting that the wheat IM is very sensitive to the activity of these genes.

Loss-of-function mutations in *TERMINAL FLOWER 1* (*TFL1*) in Arabidopsis or in the *CENTRORADIALIS* (*CEN*) homolog in *Antirrhinum* result in the formation of a terminal flower and the transformation of indeterminate into determinate inflorescences (Bradley et al., 1997; Ratcliffe et al., 1999). In rice, knockdowns of the four *CEN* homologs (*RCN1*-*RCN4*) reduced the number of branches, whereas their overexpression increased branch number by competing with rice *FT* homologs (Kaneko-Suzuki et al., 2018; Nakagawa et al., 2002). In wheat overexpression of *CEN-D2* extended the duration of the double-ridge stage and increased the number of spikelets per spike in wheat (Wang et al., 2017), whereas loss-of-function mutations in barley *CEN2* reduced the number of spikelets per spike (Bi et al., 2019). Based on these results, we hypothesize that the upregulation of the wheat *CEN2*, *CEN4* and *CEN5* homologs in the developing spike of the *vrn1ful2*-null mutant might have contributed to its indeterminate growth.

### The *vrn1*-null and *ful2*-null mutants have a higher number of spikelets per spike

We showed in this study that the timing of the transition between the IM and the terminal spikelet is modulated by *VRN1* and *FUL2* and that this affects the number of spikelets per spike. The stronger effect of *vrn1*-null (nine additional spikelets, Fig. 4C) relative to *ful2*-null (two additional spikelets, Fig. 4D) is likely associated with *VRN1*’s stronger effect on heading time (Fig. 1A-C), which provides more time for the formation of additional spikelets. This seems to be also the case in rice, where overexpression of *OsMADS15*, resulted in reducted number of primary branches in the panicle (Lu et al., 2012). Similarly, a stop codon mutation in a homolog of AP1 in rapeseed altered plant architecture and increased the number of seeds per plant (Shah et al., 2018). Taken together, these results suggest that mutations in this group of meristem identity genes may be useful to modulate seed number in different plant species.

In addition to its effect on spikelet number, the *ful2*-null mutation was also associated with an increase in the number of florets per spikelet, which suggests that this gene contributes to maintaining a limited number of florets per spikelet (Fig. S10C-F). This effect was not detected in *vrn1*-null and *ful3*-null. Since a higher number of florets per spikelet and increased spikelet number can both contribute to increases in grain yield potential, we explored the effect of the *ful2*-null mutant on grain yield components. In a field study, the *ful2*-null plants showed a 30.8% increase in the average number of grains per spike compared with the control sister lines. Although in this experiment the positive increase in grain number was partially offset by a decrease in average grain weight, the total grain weight per spike was still slightly higher (6.3%) in the *ful2*-null mutant relative to the control. It would be interesting to test if the introgression of this mutation in genotypes which high biomass (increased “source”) grown under optimum agronomic conditions can reduce the negative correlation between grain number and grain weight.

In summary, our results indicate that *VRN1*, *FUL2* and *FUL3* play redundant and essential roles in spikelet development, repression of the lower ridge and spike determinacy, and that mutations in *VRN1* and *FUL2* can be used to increase the number of spikelets per spike, an important component of grain yield. These results suggest that a better understanding of the processes that control the development of grass flowers and inflorescences may contribute to improving the productivity of a group of species that is critical for the global food supply.

## ACKNOWLEDGEMENTS

This project was supported by the Howard Hughes Medical Institute, NRI Competitive Grant 2016-67013-24617 from the USDA National Institute of Food and Agriculture (NIFA) and the International Wheat Partnership Initiative (IWYP). We thank Dr. Alejandra Alvarez and Dr. Josh Hegarty for their help with field experiments and Dr. Daniel Wood and Dr. Juan Debernardi for their valuable comments and suggestions. We thank the anonymous reviewers for their helpful comments and valuable suggestions.

## AUTHOR CONTRIBUTIONS

CL and JD designed research; JD provided overall supervision to the project; HL, CL, AC, ML and JJ performed research; CL, HL and JD analyzed data; JD provided statistical analyses; CL wrote first draft and JD the final version. All authors reviewed the paper.

## MATERIALS AND METHODS

### Selected mutations and mutant combinations

An ethyl methane sulphonate (EMS) mutagenized population of the tetraploid wheat variety Kronos was screened for mutations initially using *Cel*I assays (Uauy et al., 2009) and later using BLAST searches in the database of sequenced mutations for the same population (Krasileva et al., 2017). We identified loss-of-function mutations in the A and B genome homeologs of *FUL2* and *FUL3*, which were confirmed using genome specific primers described in Table S1. Single genome mutants were backcrossed two to three times to Kronos *vrn2*-null to reduce background mutations. The wild type Kronos carries a functional *VERNALIZATION 2* (*VRN2*) flowering repressor, which results in extremely late flowering in the presence of the *vrn1*-null mutation (Chen and Dubcovsky, 2012). To avoid this problem all mutants described in this study were developed in a Kronos *vrn2*-null background (Distelfeld et al., 2009b), unless indicated otherwise.

For *FUL-A2*, we selected line T4-837 (henceforth *ful-A2*), which has a mutation in the splice donor site of the fifth intron. RT-PCR and sequencing analysis of the *ful-A2* transcripts revealed two incorrect splicing forms. The most abundant form skipped the fifth exon, which resulted in a deletion of 14 amino acids in the middle of the K-box (Δ144-157). In the other alternative splicing form, the retention and translation of the fifth intron generated a premature stop codon that disrupted the K-box and removed the entire C-terminus (Fig. S2A). For *FUL-B2*, we selected line T4-2911 that carries a C to T change at nucleotide 484 (henceforth *ful-B2*). The *ful-B2* mutation generates a premature stop codon at position 162 (Q162*) that removed the last 13 amino acids of the K-box and the entire C-terminus (Fig. S2A).

For *FUL-A3*, we selected line T4-2375 that carries a G to A mutation in the splice acceptor site of the third intron. Sequencing of *ful-A3* transcripts revealed that this mutation generated a new splice acceptor site that shifted the reading frame by one nucleotide. The alternative translation generated a premature stop codon that truncated 72% of the K-box and the entire C-terminus (Fig. S2B). For *FUL-B3*, we selected line T4-2139 that carries a C to T mutation at nucleotide position 394 that generates a premature stop codon at amino acid position 132 (Q132*). This premature stop removed half of the K-box and the complete C-terminus (Fig. S2B). Given the critical roles of the K-domain in protein-protein interactions, and the C-terminal domain in transcriptional activation, these selected mutations are expected to impair the normal function of the FUL2 and FUL3 proteins.

The A and B-genome mutants for each gene were intercrossed to generate double mutants, which for simplicity, are referred to hereafter as null mutants. The *ful2*-null and *ful3*-null were intercrossed with a *vrn1vrn2*-null mutant (*vrn-A1*-null T4-2268 / *vrn-B1* T4-2619 / *vrn2*-null) (Chen and Dubcovsky, 2012) to generate *vrn1ful2*-null and *vrn1ful3*-null, which were finally intercrossed to generate *vrn1ful2ful3*-null (all in a *vrn2*-null background). The *vrn1ful2*-null and *vrn1ful2ful3*-null mutants were sterile, so they were maintained and crossed by keeping the *ful-B2* mutation in heterozygous state. The single mutants are available as part of the public sequenced TILLING populations (Krasileva et al., 2017) and the mutant combinations are available upon request.

Transgenic Kronos plants overexpressing *FUL2* and *FUL3* coding regions were generated at the UC Davis Transformation facility using *Agrobacterium*-mediated transformation. The coding regions of these two genes were cloned from Kronos into the binary vector pLC41 (Japan Tobacco, Tokyo, Japan) downstream of the maize *UBIQUITIN* promoter. A C-terminal 3xHA tag was added to FUL2 and a C-terminal 4xMYC tag was added to FUL3. Mutant and transgenic wheat plants were grown in PGR15 CONVIRON chambers under LD (16h light/8h dark, light intensity ~330 μM m^−2^ s^−1^) at 22 °C during the day and 18 °C during the night.

To study the effect of *Ubi::FUL2* in different genetic backgrounds we crossed the Kronos *Ubi::FUL2* with Kronos-*vrn1vrn2*-null and analyzed the effect of the three genes in the F_2_ progeny under greenhouse conditions. A field experiment comparing *ful2*-null and its control line was performed at the University of California, Davis field station during the 2017-2018 growing season (sowed on 12/1/2017 and harvested on 06/25/2018). One meter rows (30 grains per row) were used as experimental units and the experiment was organized in a randomized complete block design with eight blocks. During the growing season plants received 200 units of N as ammonium sulfate, three irrigations, one application of broad-leaf herbicides (2, 4D + Buctril) and alternating applications of fungicides Quadris and Tilt every 2 weeks.

### Effect of the splice site mutations in *ful-A2* and *ful-B3* mutants

To determine the effect of the splice site mutations in *ful-A2* and *ful-B3*, we extracted total RNA from leaf samples using the Spectrum™ Plant Total RNA kit. cDNA was synthesized from 2 μg of RNA using the High Capacity Reverse Transcription Kit according to the manufacturer’s instructions and used as RT-PCR template. For *ful-A2*, we used primers *FUL2-837-F* (5’-CCATACAAAAATGTCACAAGC-3’) and *FUL2-837-R* (5’-TTCTGC CTCTCCACCAGTT-3’) for RT-PCR. These primers, which are not genome specific, amplified three fragments of 303 bp, 220 bp and 178 bp. We gel-purified these fragments, cloned them into pGEM-T vector (Promega), and sequenced them. The 220 bp fragment was from the wild type *FUL-B2* allele, whereas the other two fragments corresponded to two alternative splicing forms of *ful-A2* that either retained the fifth intron (303 bp) or skipped the fifth exon (178 bp).

For the *ful-B3* mutant, we performed RT-PCR using primers FUL3-2375-F (5’-ATGGATGTGATTCTTGAAC-3’) and FUL3-2375-R (5’-TGTCCTGCAGAAGCACCTCGTAGAGA-3’). Sequencing analysis of the PCR products showed that the G to A mutation generated a new splice acceptor site with an adjacent G that shifted the reading frame by one nucleotide after 333 bp, and generated a premature stop codon.

### Scanning Electron-Microscopy (SEM)

Apices from different developmental stages were dissected and fixed for a minimum of 24 h in FAA (50% ethanol, 5% (v/v) acetic acid, 3.7% (v/v) formaldehyde), and then dehydrated through a graded ethanol series to absolute ethanol. Samples were critical-point dried in liquid CO2 (tousimis ^®^ 931 Series critical point drier), mounted on aluminum stubs, sputter-coated with gold (Bio-Rad SEM Coating System Model E5100), and examined with a Philips XL30 scanning electron-microscope operating at 5KV. Images were recorded at slow scan 3 for high definition and saved as TIFF files.

### RNA extraction and Real-time qPCR analysis

RNAs from apices were extracted using the Trizol reagent (ThermoFisher Scientific, Cat. No.15596026). One μg of RNA was used for cDNA synthesis following the instructions of the “High-Capacity cDNA Reverse Transcription Kit” (ThermoFisher Scientific, Cat. No. 4368814). The cDNA was then diluted 20 times and 5 μl of the dilution was mixed with 2×VeriQuest Fast SYBR Green qPCR Master Mix (Affymetrix, Cat. No. 75690) and with primers for the real-time qPCR analysis. Primer sequences are listed in Table S2. *INNITIATION FACTOR 4A* (*IF4A*) and *ACTIN* were used as an endogenous controls. Transcript levels for all genes are expressed as linearized fold-*IF4A* levels calculated by the formula 2^(*IF4A* C_T_ – *TARGET* C_T_)^ ± standard error (SE) of the mean. The resulting number indicates the ratio between the initial number of molecules of the target gene and the number of molecules in the endogenous control.

## SUPPLEMENTARY TABLES

**Table S1.** Primers used for genotyping the *ful2*, *ful3, vrn1* and *vrn2*-null mutants.

**Table S2.** Primers used in the real time Q-PCR experiments.

**Table S3.** Three-way ANOVA for heading time.

## SUPPLEMENTARY FIGURES

**Fig. S1.** Phylogeny of duplicated Arabidopsis AP1/CAL/FUL and grasses VRN1/FUL2/FUL3 proteins.

**Fig. S2**. Selected *ful2* and *ful3* mutations and their effect on the encoded proteins.

**Fig. S3.** Time course of shoot apical meristem (SAM) elongation and transition to the reproductive stage in *FUL2* and *FUL3* mutants in a *vrn1*-null background.

**Fig. S4.** Effect of *Ubi::FUL2* and *Ubi::FUL3* on heading time.

**Fig. S5.** *FT1* transcript levels in leaves.

**Fig. S6.** Phenotypical characterization of heterozygous mutants containing one copy of *VRN1* or *FUL2*.

**Fig. S7.** Transcript levels of wheat *SVP*-like MADS-box genes *VRT2*, *BM1* and *BM10*.

**Fig. S8**. Effect of the overexpression of *FUL2* and *FUL3* on the number of spikelets per spike.

**Fig. S9**. Transcript levels of wheat *TFL1/CEN*-like genes *CEN2*, *CEN4* and *CEN5*.

**Fig. S10**. Effect of *ful2*-null and *ful3*-null on the number of florets per spikelet.

## REFERENCES

Alonso-Peral, M. M., Oliver, S. N., Casao, M. C., Greenup, A. A. and Trevaskis, B. (2011). The promoter of the cereal *VERNALIZATION1* gene is sufficient for transcriptional induction by prolonged cold. PLoS One 6, e29456.

Bemer, M., van Mourik, H., Muino, J. M., Ferrandiz, C., Kaufmann, K. and Angenent, G. C. (2017). FRUITFULL controls *SAUR10* expression and regulates Arabidopsis growth and architecture. Journal of Experimental Botany 68, 3391–3403.

Bi, X., van Esse, G. W., Mulki, M. A., Kirschner, G., Zhong, J., Simon, R. and von Korff, M. (2019). *CENTRORADIALIS* interacts with *FLOWERING LOCUS T*-like genes to control floret development and grain number. Plant Physiology On line first., DOI:10.1104/pp.18.01454.

Bradley, D., Ratcliffe, O., Vincent, C., Carpenter, R. and Coen, E. (1997). Inflorescence commitment and architecture in Arabidopsis. Science 275, 80–83.

Chen, A. and Dubcovsky, J. (2012). Wheat TILLING mutants show that the vernalization gene *VRN1* down-regulates the flowering repressor *VRN2* in leaves but is not essential for flowering. PLoS Genetics 8, e1003134.

Ciaffi, M., Paolacci, A. R., Tanzarella, O. A. and Porceddu, E. (2011). Molecular aspects of flower development in grasses. Sexual Plant Reproduction 24, 247–282.

Debernardi, J. M., Lin, H., Chuck, G., Faris, J. D. and Dubcovsky, J. (2017). microRNA172 plays a crucial role in wheat spike morphogenesis and grain threshability. Development 144, 1966–1975.

Distelfeld, A., Li, C. and Dubcovsky, J. (2009a). Regulation of flowering in temperate cereals. Current Opinion in Plant Biology 12, 178–184.

Distelfeld, A., Tranquilli, G., Li, C., Yan, L. and Dubcovsky, J. (2009b). Genetic and molecular characterization of the *VRN2* loci in tetraploid wheat. Plant Physiology 149, 245–257.

Ferrándiz, C., Gu, Q., Martienssen, R. and Yanofsky, M. F. (2000). Redundant regulation of meristem identity and plant architecture by *FRUITFULL*, *APETALA1* and *CAULIFLOWER*. Development 127, 725–734.

Friend, D. J. C. (1965). Tillering and leaf production in wheat as affected by temperature and light intensity. Canadian Journal of Botany 43, 1063–&.

Fu, D., Szűcs, P., Yan, L., Helguera, M., Skinner, J., Hayes, P. and Dubcovsky, J. (2005). Large deletions within the first intron in *VRN-1* are associated with spring growth habit in barley and wheat. Molecular Genetics and Genomics 273, 54–65.

Gocal, G. F. W., King, R. W., Blundell, C. A., Schwartz, O. M., Andersen, C. H. and Weigel, D. (2001). Evolution of floral meristem identity genes. Analysis of *Lolium temulentum* genes related to *APETALA1* and *LEAFY* of Arabidopsis. Plant Physiology 125, 1788–1801.

Gómez-Ariza, J., Brambilla, V., Vicentini, G., Landini, M., Cerise, M., Carrera, E., Shrestha, R., Chiozzotto, R., Galbiati, F., Caporali, E. et al. (2019). A transcription factor coordinating internode elongation and photoperiodic signals in rice. Nature Plants 5, 358–362.

Honma, T. and Goto, K. (2001). Complexes of MADS-box proteins are sufficient to convert leaves into floral organs. Nature 409, 525–9.

Kaneko-Suzuki, M., Kurihara-Ishikawa, R., Okushita-Terakawa, C., Kojima, C., Nagano-Fujiwara, M., Ohki, I., Tsuji, H., Shimamoto, K. and Taoka, K. I. (2018). TFL1-like proteins in rice antagonize rice FT-like protein in inflorescence development by competition for complex formation with 14-3-3 and FD. Plant and Cell Physiology 59, 458–468.

Kaufmann, K., Wellmer, F., Muino, J. M., Ferrier, T., Wuest, S. E., Kumar, V., Serrano-Mislata, A., Madueno, F., Krajewski, P., Meyerowitz, E. M. et al. (2010). Orchestration of floral initiation by APETALA1. Science 328, 85–89.

Kellogg, E. A. (2001). Evolutionary history of the grasses. Plant Physiology 125, 198–1205.

Kellogg, E. A., Camara, P. E. A. S., Rudall, P. J., Ladd, P., Malcomber, S. T., Whipple, C. J. and Doust, A. N. (2013). Early inflorescence development in the grasses (Poaceae). Frontiers in Plant Science 4, 250.

Kippes, N., Chen, A., Zhang, X., Lukaszewski, A. J. and Dubcovsky, J. (2016). Development and characterization of a spring hexaploid wheat line with no functional *VRN2* genes. Theoretical and Applied Genetics 129, 1417–1428.

Kobayashi, K., Yasuno, N., Sato, Y., Yoda, M., Yamazaki, R., Kimizu, M., Yoshida, H., Nagamura, Y. and Kyozuka, J. (2012). Inflorescence meristem identity in rice is specified by overlapping functions of three *AP1/FUL*-like MADS box genes and *PAP2*, a *SEPALLATA* MADS box gene. Plant Cell 24, 1848–1859.

Krasileva, K. V., Vasquez-Gross, H., Howell, T., Bailey, P., Paraiso, F., Clissold, L., Simmonds, J., Ramirez-Gonzalez, R. H., Wang, X., Borrill, P. et al. (2017). Uncovering hidden variation in polyploid wheat. Proceedings of the National Academy of Sciences of the United States of America 114, E913–E921

Li, Q., Wang, Y., Wang, F. X., Guo, Y. Y., Duan, X. Q., Sun, J. H. and An, H. L. (2016). Functional conservation and diversification of *APETALA1/FRUITFULL* genes in *Brachypodium distachyon*. Physiologia Plantarum 157, 507–518.

Liu, C., Zhou, J., Bracha-Drori, K., Yalovsky, S., Ito, T. and Yu, H. (2007). Specification of Arabidopsis floral meristem identity by repression of flowering time genes. Development 134, 1901–1910.

Lu, S. J., Wei, H., Wang, Y., Wang, H. M., Yang, R. F., Zhang, X. B. and Tu, J. M. (2012). Overexpression of a transcription factor *OsMADS15* modifies plant architecture and flowering time in rice (*Oryza sativa* L.). Plant Molecular Biology Reporter 30, 1461–1469.

Lv, B., Nitcher, R., Han, X., Wang, S., Ni, F., Li, K., Pearce, S., Wu, J., Dubcovsky, J. and Fu, D. (2014). Characterization of *FLOWERING LOCUS T1* (*FT1*) gene in *Brachypodium* and wheat. PLoS One 9, e94171.

Malcomber, S. T., Preston, J. C., Reinheimer, R., Kossuth, J. and Kellogg, E. A. (2006). Developmental gene evolution and the origin of grass inflorescence diversity. Advances in Botanical Research 44, 425–481.

McSteen, P., Laudencia-Chingcuanco, D. and Colasanti, J. (2000). A floret by any other name: control of meristem identity in maize. Trends in Plant Science 5, 61–66.

Nakagawa, M., Shimamoto, K. and Kyozuka, J. (2002). Overexpression of *RCN1* and *RCN2*, rice *TERMINAL FLOWER 1/CENTRORADIALIS* homologs, confers delay of phase transition and altered panicle morphology in rice. Plant Journal 29, 743–750.

Oikawa, T. and Kyozuka, J. (2009). Two-step regulation of LAX PANICLE1 protein accumulation in axillary meristem formation in rice. Plant Cell 21, 1095–1108.

Pearce, S., Vanzetti, L. S. and Dubcovsky, J. (2013). Exogenous gibberellins induce wheat spike development under short days only in the presence of *VERNALIZATION 1*. Plant Physiology 163, 1433–1445.

Preston, J. C., Christensen, A., Malcomber, S. T. and Kellogg, E. A. (2009). MADS-box gene expression and implications for developmental origins of the grass spikelet. American Journal of Botany 96, 1419–1429.

Preston, J. C. and Kellogg, E. A. (2006). Reconstructing the evolutionary history of paralogous *APETALA1/FRUITFULL*-like genes in grasses (Poaceae). Genetics 174, 421–437.

Preston, J. C. and Kellogg, E. A. (2008). Discrete developmental roles for temperate cereal grass *VRN1*/*FUL*-like genes in flowering competency and the transition to flowering. Plant Physiology 146, 265–276.

Ratcliffe, O. J., Bradley, D. J. and Coen, E. S. (1999). Separation of shoot and floral identity in Arabidopsis. Development 126, 1109–1120.

Ream, T. S., Woods, D. P., Schwartz, C. J., Sanabria, C. P., Mahoy, J. A., Walters, E. M., Kaeppler, H. F. and Amasino, R. M. (2014). Interaction of photoperiod and vernalization determines flowering time of *Brachypodium distachyon*. Plant Physiology 164, 694–709.

Shah, S., Karunarathna, N. L., Jung, C. and Emrani, N. (2018). An *APETALA1* ortholog affects plant architecture and seed yield component in oilseed rape (*Brassica napus* L.). BMC Plant Biol 18, 380.

Shaw, L. M., Lyu, B., Turner, R., Li, C., Chen, F., Han, X., Fu, D. and Dubcovsky, J. (2019). *FLOWERING LOCUS T2* (*FT2*) regulates spike development and fertility in temperate cereals. Journal of Experimental Botany 70, 193–204.

Theissen, G., Melzer, R. and Rumpler, F. (2016). MADS-domain transcription factors and the floral quartet model of flower development: linking plant development and evolution. Development 143, 3259–3271.

Tranquilli, G. and Dubcovsky, J. (2000). Epistatic interactions between vernalization genes *Vrn-A^m^1* and *Vrn-A^m^2* in diploid wheat. Journal of Heredity 91, 304–306.

Trevaskis, B., Tadege, M., Hemming, M. N., Peacock, W. J., Dennis, E. S. and Sheldon, C. (2007). *Short Vegetative Phase-like* MADS-box genes inhibit floral meristem identity in barley. Plant Physiology 143, 225–235.

Uauy, C., Paraiso, F., Colasuonno, P., Tran, R. K., Tsai, H., Berardi, S., Comai, L. and Dubcovsky, J. (2009). A modified TILLING approach to detect induced mutations in tetraploid and hexaploid wheat. BMC Plant Biology 9, 115–128.

Wang, Y. G., Yu, H. P., Tian, C. H., Sajjad, M., Gao, C. C., Tong, Y. P., Wang, X. F. and Jiao, Y. L. (2017). Transcriptome association identifies regulators of wheat spike architecture. Plant Physiology 175, 746–757.

Whipple, C. J. (2017). Grass inflorescence architecture and evolution: the origin of novel signaling centers. New Phytologist 216, 367–372.

Williams, R. F. and Langer, R. H. M. (1975). Growth and development of the wheat tiller. II. The dynamics of tiller growth. Australian Journal of Botany 23, 745–759.

Woods, D. P., McKeown, M. A., Dong, Y. X., Preston, J. C. and Amasino, R. M. (2016). Evolution of *VRN2/Ghd7*-like genes in vernalization-mediated repression of grass flowering. Plant Physiology 170, 2124–2135.

Wu, F., Shi, X. W., Lin, X. L., Liu, Y., Chong, K., Theissen, G. and Meng, Z. (2017). The ABCs of flower development: mutational analysis of *AP1/FUL*-like genes in rice provides evidence for a homeotic (A)-function in grasses. Plant Journal 89, 310–324.

Yan, L., Fu, D., Li, C., Blechl, A., Tranquilli, G., Bonafede, M., Sanchez, A., Valarik, M. and Dubcovsky, J. (2006). The wheat and barley vernalization gene *VRN3* is an orthologue of *FT*. Proceedings of the National Academy of Sciences of the United States of America 103, 19581–19586.

Yan, L., Loukoianov, A., Tranquilli, G., Helguera, M., Fahima, T. and Dubcovsky, J. (2003). Positional cloning of wheat vernalization gene *VRN1*. Proceedings of the National Academy of Sciences of the United States of America 100, 6263–6268.

Zhang, X. K., Xia, X. C., Xiao, Y. G., Dubcovsky, J. and He, Z. H. (2008). Allelic variation at the vernalization genes *Vrn-A1*, *Vrn-B1*, *Vrn-D1* and *Vrn-B3* in Chinese common wheat cultivars and their association with growth habit. Crop Science 48, 458–470.

