## Supplementary Tables and Figures for "Wheat *VRN1*, *FUL2* and *FUL3* play critical and redundant roles in spikelet development and spike determinacy"

### Supplementary Tables

**Table S1**. Primers used for genotyping the *ful2*, *ful3, vrn1* and *vrn2*-null mutants

| **Mutant** | | **Primer sequences** | **Marker type and restriction enzyme** |
|---|---|---|---|
| ***ful-3A*** | Forward | ATGGATGTGATTCTTGAAC | CAPS (*Sml*I) |
| | Reverse | AAACGTAAATACAGTGGAAC | |
| ***ful-3B*** | Forward | GAAGAGTCAAAGGTATTGTATTTCTA | dCAPS (*Pvu*II) |
| | Reverse | TGCTTCAAAGAACTATCCAGCT | |
| ***ful-2A*** | Forward | CACATTACCTTACCATCTACTCAC | CAPS (*HpyA*V) |
| | Reverse | CATGCAACAGAATTATACAGC | |
| ***ful-2B*** | Forward | CCATTGACCATCAACTAGATCA | CAPS (*Btsα*I) |
| | Reverse | TGAACCAGCATTCCTGATA | |
| ***vrn-1A*** | Forward | GCCTATTTGTAGCATTTCTGTCATT | CAPS (*Bsg*I) [1] |
| | Reverse | GAACATCTCAGTCTAGAATCTGAT | |
| ***vrn-1B*** | Forward | CGCCTCACCCAACCACCTGAC | CAPS (*Bsl*I) [1] |
| | Reverse | ACGGATGGAAACAGCTACCGA | |
| ***vrn2*** | Forward | AACGCTTTATGATGCCAAGG | CAPS for linked *SNF2*(*HpyCh4*IV) [2] |
| | Reverse | TGTGGACAGAACTGGTTTGC | |

[1] **Chen, A. and Dubcovsky, J.** (2012). *PLoS Genetics* **8**, e1003134.

[2] **Distelfeld, A., Tranquilli, G., Li, C., Yan, L. and Dubcovsky, J.** (2009). *Plant Physiology* **149**, 245-257.

**Table S2**. Primers used the real time Q-PCR experiments.

| Gene | | Primer sequence | Primer efficiency |
|---|---|---|---|
| ***VRT2*** | Forward | AGCAGCCGGCTATTGATTTA | 101% |
| | Reverse | TTCCATCTGCTGCAACTCAC | |
| ***BM1*** | Forward | CCAGCAGATGAGAGGAGAGG | 100% |
| | Reverse | GGCTCTTGGTTTTCAGAACG | |
| ***BM10*** | Forward | CACAGGGTGCTTCAGACAAA | 103% |
| | Reverse | TTGGAATCTGGCCTACTTGG | |
| ***CEN-2*** | Forward | CTCGTCGTTGGAAAGGTCAT | 105% |
| | Reverse | TGAACACCTGCTTGTTGGAG | |
| ***CEN-4*** | Forward | TCGTCGTCGGAAAGGTCATC | 100% |
| | Reverse | GAACTCCTGGCCATTGGACA | |
| ***CEN-5*** | Forward | GGCCATGAGCTCTACCCATC | 107% |
| | Reverse | ACCCCCTTGGACCTCTACTC | |
| ***FT1*** | Forward | CAGCAGCCCAGGGTTGAG | 100% [1] |
| | Reverse | ATCTGGGTCTACCATCACGAGTG | |

[1] **Yan L, Fu D, Li C, Blechl A, Tranquilli G, Bonafede M, Sanchez A, Valarik M, Yasuda S, Dubcovsky J.** (2006) PNAS **103**:19581-6.

**Table S3**. Three-way ANOVA for heading time. Population segregating for *VRN1* (*vrn1*-null vs. heterozygous *Vrn1*), *VRN2* (*vrn2*-null vs. wild type *Vrn2*) and *FUL2* (*Ubi::FUL2* versus non-transgenic control). The population was generated from the cross between Kronos *vrn1vrn2*-null and Kronos overexpressing *FUL2* under the maize *UBIQUITIN* promoter (*Vrn1Vrn2*). Normality of residuals was confirmed by the Shapiro-Wilk test ($P$ = 0.3516).

Dependent Variable: HD

| Source | DF | Sum of Squares | Mean Square | F Value | Pr > F |
|---|---|---|---|---|---|
| Model | 7 | 19506.2 | 2786.6 | 311.93 | <.0001 |
| Error | 47 | 419.9 | 8.9 | | |
| Corrected Total | 54 | 19926.1 | | | |

$R^2$= 0.978929

| Source | DF | Type III SS | Mean Sq. | F Value | Pr > F |
|---|---|---|---|---|---|
| VRN1 | 1 | 7339.3 | 7339.3 | 821.57 | <.0001 |
| FUL2 | 1 | 3336.2 | 3336.2 | 373.46 | <.0001 |
| VRN1*FUL2 | 1 | 1502.7 | 1502.7 | 168.21 | <.0001 |
| VRN2 | 1 | 13441.2 | 13441.2 | 1504.61 | <.0001 |
| VRN1*VRN2 | 1 | 2366.2 | 2366.2 | 264.88 | <.0001 |
| FUL2*VRN2 | 1 | 1696.9 | 1696.9 | 189.95 | <.0001 |
| VRN1*FUL2*VRN2 | 1 | 717.9 | 717.9 | 80.36 | <.0001 |

### Supplementary Figures

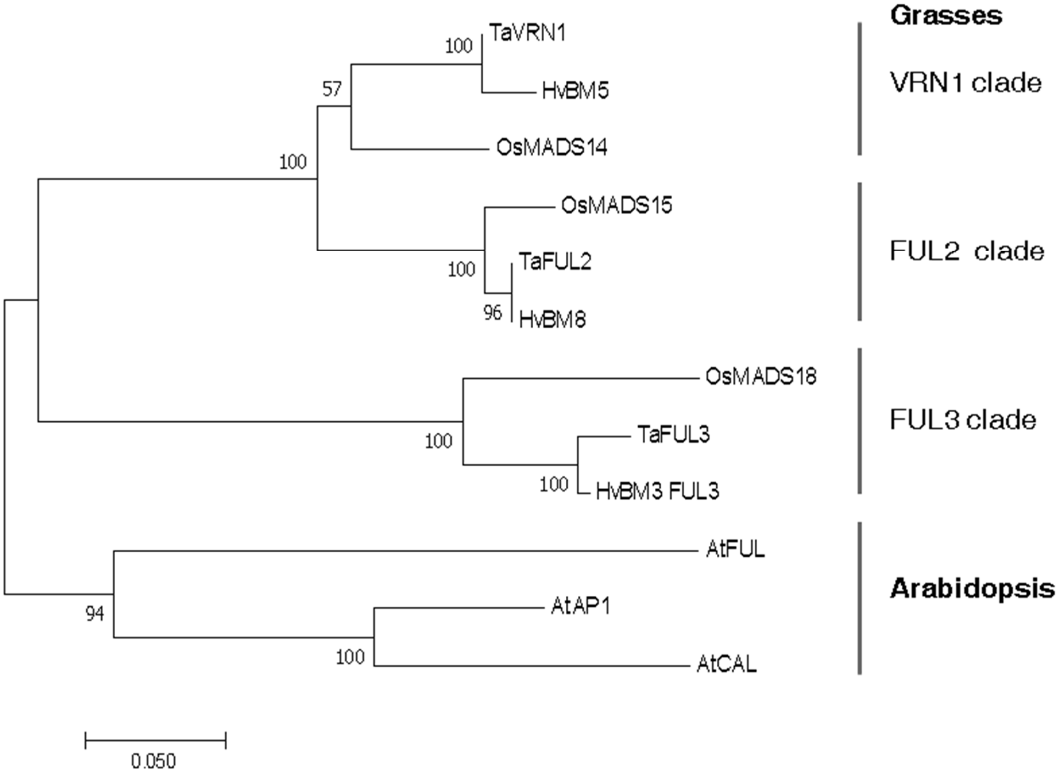


**Fig. S1. Phylogeny of duplicated Arabidopsis AP1/CAL/FUL and grasses VRN1/FUL2/FUL3 clades.** Evolutionary history inferred using the Neighbor-Joining method (Saitu and Nei, 1987). The optimal tree is shown with bootstrap values >50 shown in the nodes (based on 1000 replicates, Felsenstein, 1985). The tree is drawn to scale, with branch lengths in the same units as those of the evolutionary distances used to infer the phylogenetic tree. The evolutionary distances were computed using the Poisson correction method (Zuckerkandl and Pauling, 1965) and are in the units of the number of amino acid substitutions per site. All positions containing gaps and missing data were eliminated. There were a total of 176 positions in the final dataset. Evolutionary analyses were conducted in MEGA7 (Kumar et al. 2016). ). The duplication that originated the VRN1 and FUL2 clusters occurred close to the base of the grass divergence, whereas the duplication that originated the FUL3 clade occurred close to the base of the monocots (Preston and Kellogg, 2006). At = *Arabidopsis thaliana*, Os = *Oryza sativa*, Hv = *Hordeum vulgare*, Ta = *Triticum aestivum.*

**References for Fig. S1.**


**Felsenstein J.** (1985). *Evolution* **39**,783-791.

**Kumar S., Stecher G., and Tamura K**. (2016). *Molecular Biology and Evolution 33*, 1870-1874.

**Preston, J. C. and Kellogg, E. A.** (2006). *Genetics* **174**, 421-437.

**Saitou N. and Nei M.** (1987). *Molecular Biology and Evolution* **4**,406-425.

**Zuckerkandl E. and Pauling L.** (1965). Ed. by V. Bryson and H.J. Vogel, pp. 97-166. Acad. Press, New York.

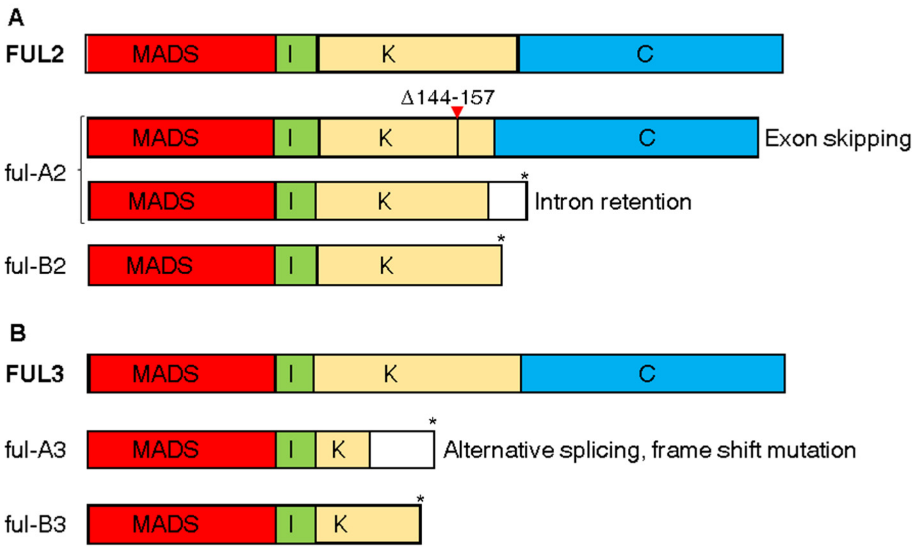


**Fig. S2. Selected *ful2* and *ful3* mutations and their effect on the encoded proteins.** Wild type FUL2 and FUL3 MADS-box proteins showing conserved domains are included as reference. MADS= MADS box domain, I= Intervening domain, K = Keratin-like box domain, C = C-terminal domain. (A) *ful-A2* mutation in the splice donor site of the 5$^{th}$ intron results in two alternative splice forms. The first one skips the 5$^{th}$ exon generating a deletion of 14 amino acids in the K-box (Δ144-157) and the second one retains the 5$^{th}$ intron resulting in altered translation (white rectangle) and a premature stop codon. The *ful-B2* mutation generates a premature stop codon (Q162*). (B) The *ful-A3* mutation in the splice acceptor site of the 3$^{rd}$ intron shifts the reading frame by one nucleotide generating a premature stop codon. The *ful-B3* mutation generates a premature stop codon (Q132*). More detailed descriptions are presented in Material and Methods.

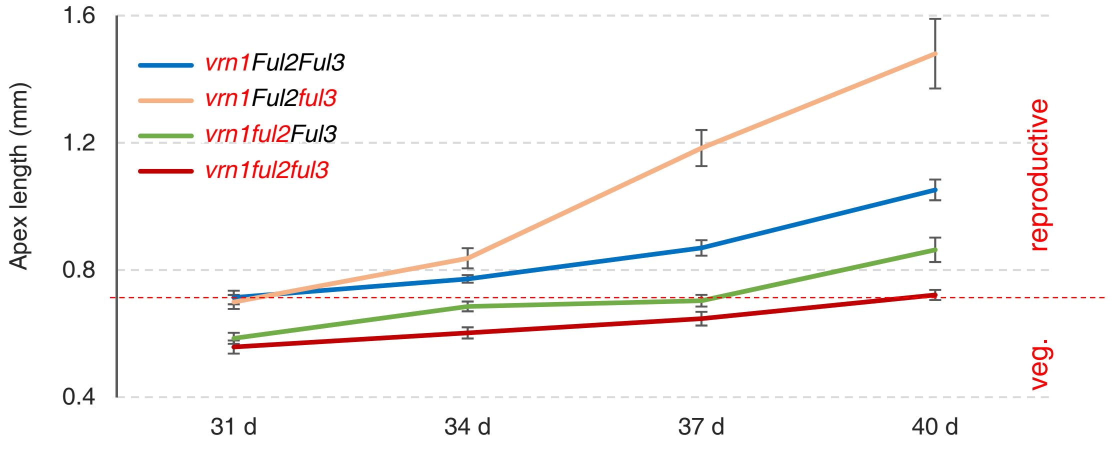


**Fig. S3. Time course of shoot apical meristem (SAM) elongation and transition to the reproductive stage in *FUL2* and *FUL3* mutants in a *vrn1*-null background.** The length of the SAM was measured every 3 d from day 31 when we observed elongation in the *vrn1Ful2Ful3* control. The red dotted line indicates the transition of the SAM to the reproductive stage. Six SAMs were measured per time point genotype combination. Error bars are standard errors of the means.

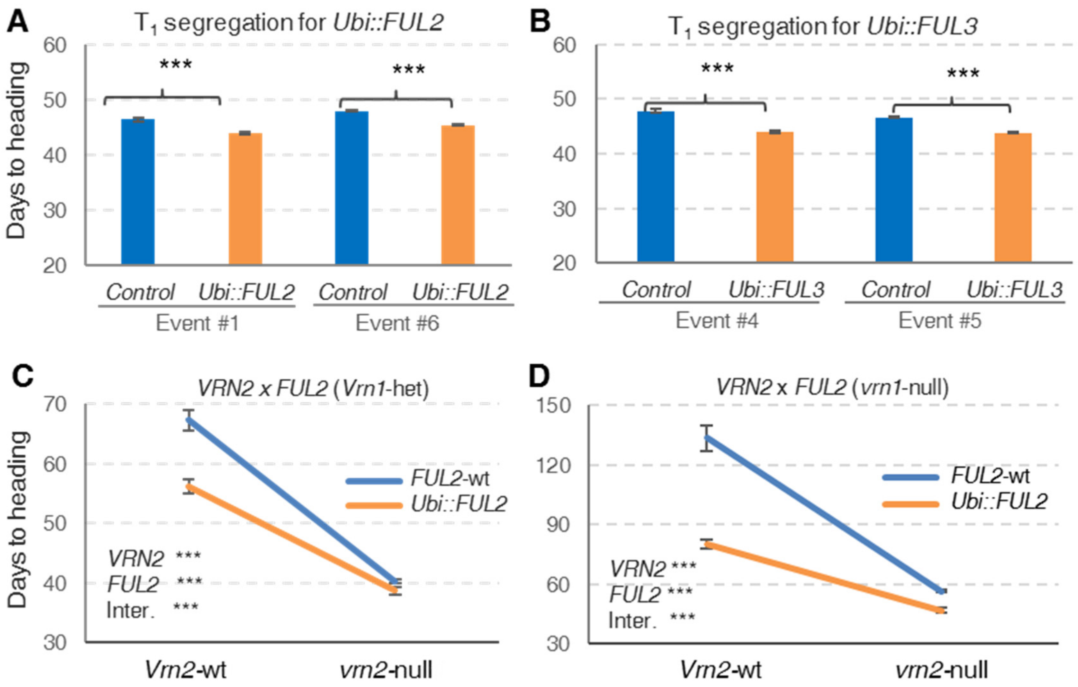


**Fig. S4**. **Effect of *Ubi::FUL2* and *Ubi::FUL3* on heading time**. (A-B) Heading time of Kronos $T_1$ transgenic plants from two independent events segregating for (A) *FUL2* (*Ubi::FUL2*, n= 4-23) and (B) *FUL3* (*Ubi::FUL3*, n= 9-31). (C-D) Two way interactions for $F_2$ plants segregating for *VRN1, VRN2* and *FUL2*. (C) *VRN2 x FUL2* in a *Vrn1* heterozygous background. (D) *VRN2 x FUL2* in a *vrn1*-null background. *P* values correspond to a 2 x 2 factorial ANOVA (3-way ANOVA in Table S4). Error bars are SEM. *** = $P < 0.0001$, NS = $P > 0.05$.

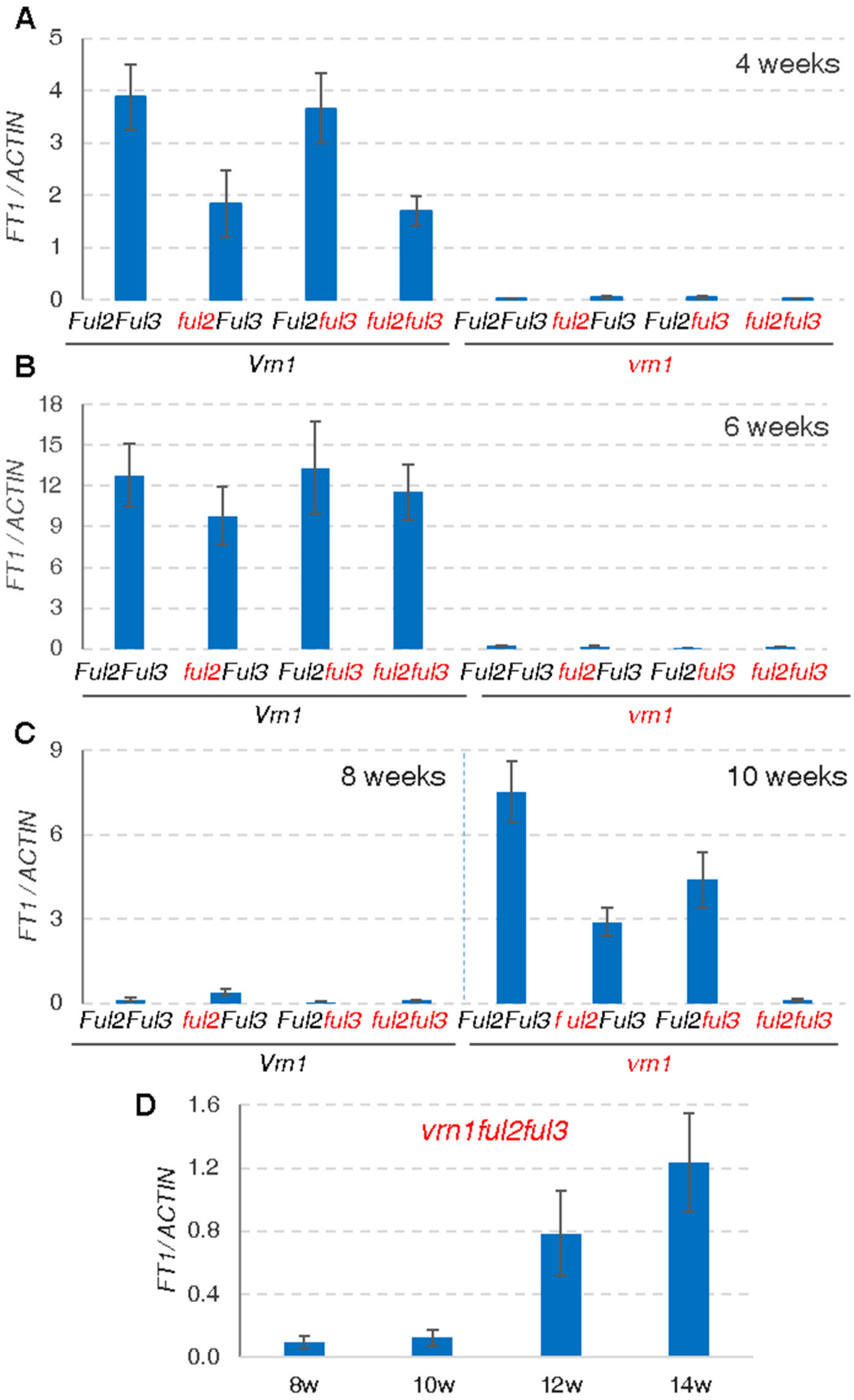


**Fig. S5. *FT1* transcript levels in leaves**. (A-B) Eight homozygous combinations of *VRN1*, *FUL2* and *FUL3* (mutant alleles in red). (A) 4-week old plants. (C) 8- and 10-week old plants (only *vrn1*-null genotypes). (D) Triple *vrn1ful2ful3*-null mutant 8- to 14-week old plants. All these mutants are in a Kronos *vrn2*-null background. Transcript levels were calculated relative to the *ACTIN* as endogenous control using the ΔCt method (scales are comparable across graphs).

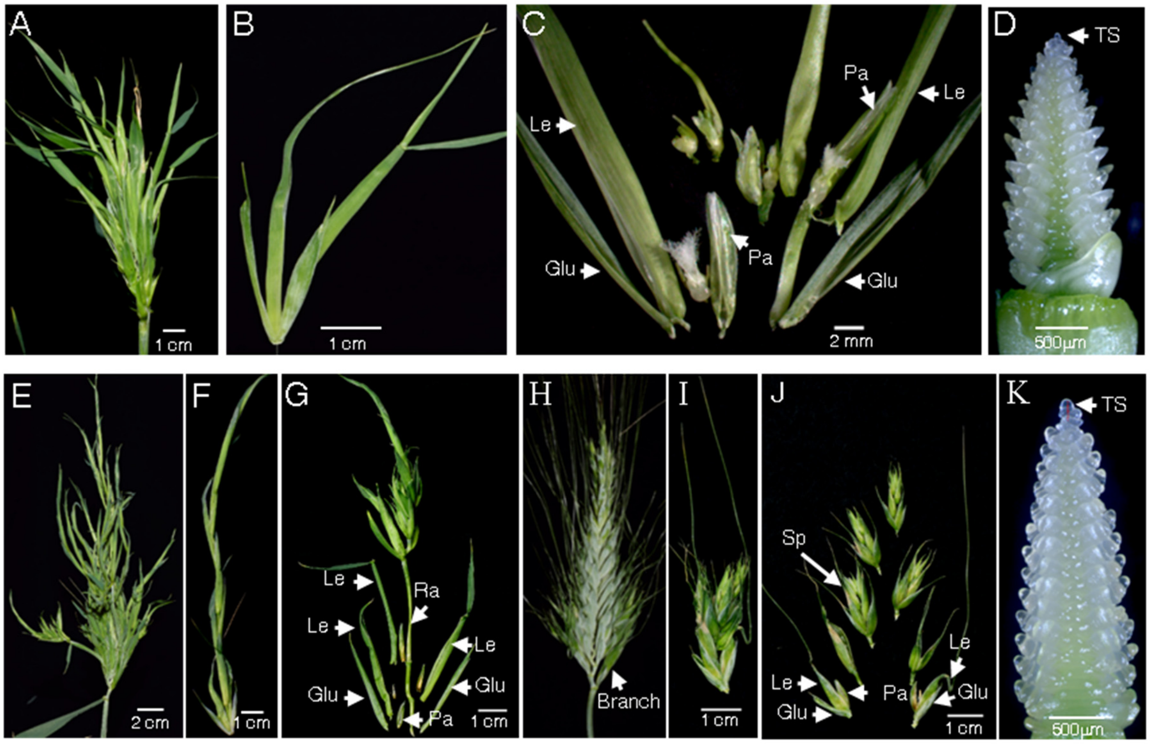


**Fig. S6**. **Phenotypical characterization of heterozygous mutants containing one copy of *VRN1* or *FUL2***. (A-C) *ful2*-null/<u>*Vrn-A1*</u> *vrn-B1*-null. (A) "Head" at early stage. (B) Detached spikelet. (C) Dissection of an individual spikelet showing floral organs including lodicules, ovaries and stamens. (D-G) *ful2*-null/*vrn-A1*-null <u>*vrn-B1*</u>. (D) Terminal spikelet (TS) confirming spike meristem determinacy. (E) Head at later stage. (F) Detached "spikelet" showing indeterminate growth. (G) Spikelet dissection showing rachilla elongation. (H-K) *vrn1*-null/*ful2-A* <u>*Ful2-B*</u> (produces viable grain). (H) Representative spikes showing formation of lateral branches in the basal region of the spike and extra florets in the terminal spikelet. (I) Detached lateral branch. (J) Dissection of the lateral branch. (K) Terminal spikelet (TS) confirming spike meristem determinacy. Sp= spikelets, Glu= glumes, Pa= palea, Le= lemma, Ra= rachilla. All these mutants are in a Kronos *vrn2*-null background.

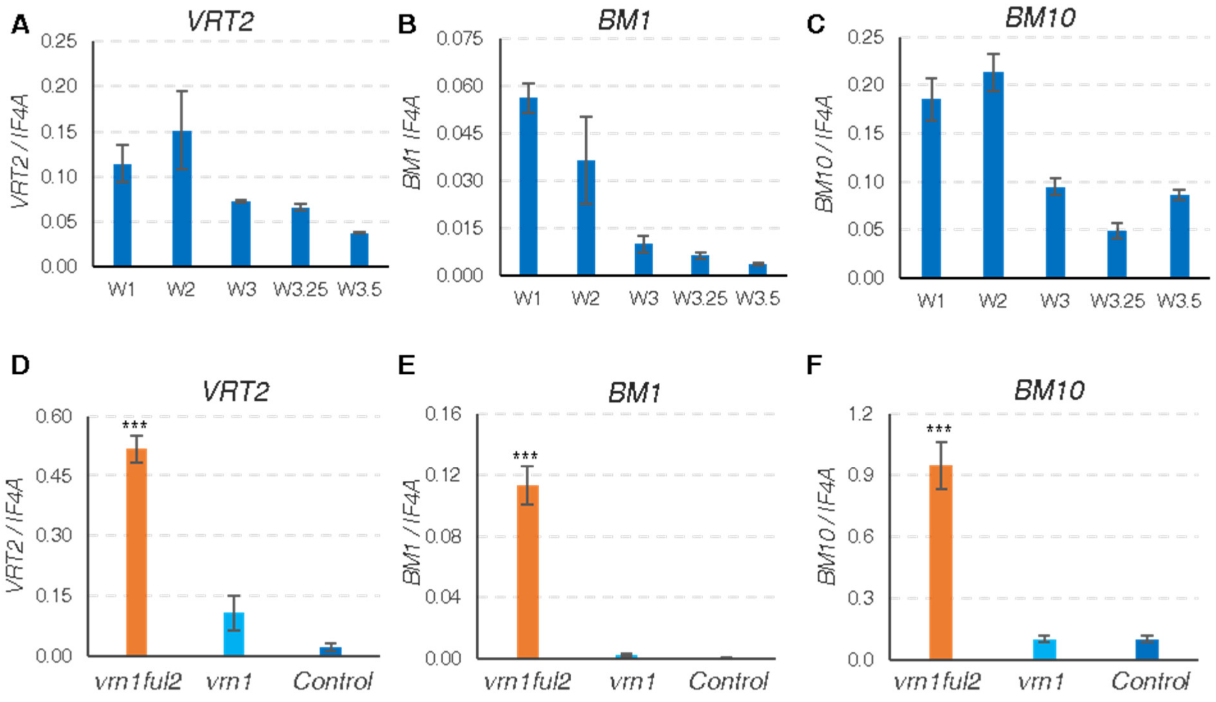


**Fig. S7. Transcript levels of wheat *SVP*-like MADS-box genes *VRT2*, *BM1* and *BM10.*** Transcript levels of (A & D) *VRT2* (*TraesCS7A01G175200* and *TraesCS7B01G080300*), (B & E) *BM1* (*TraesCS4B01G302600*), (C & F) *BM10* (*TraesCS6A01G313800* and *TraesCS6B01G343900*). (A-C) Normal spike development in wild type Kronos *(Vrn1Vrn2*) using the Waddington scale (W1.0= vegetative stage, W2.0 = double ridge stage, W3.0 = glume primordium present, W3.25 = lemma primordium present, W3.5 = and floret primordium present). (D-F) *vrn1ful2*-null, *vrn1*-null, and control at W3.5 (all in *vrn2*-null background). Transcript levels were calculated relative to the *INITIATION FACTOR 4A* (*IF4A*) as endogenous control using the ΔCt method. *** = $P < 0.0001$.

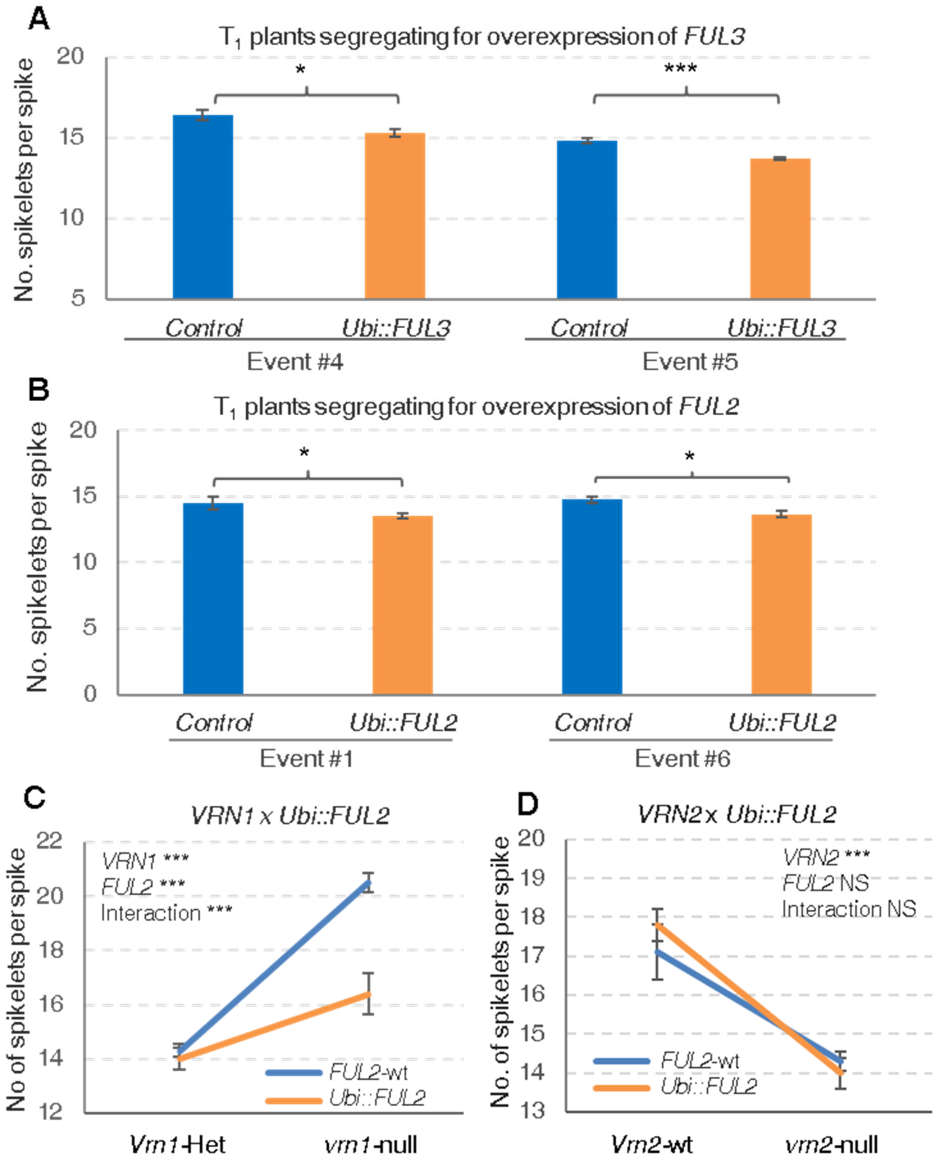


**Fig. S8. Effect of the overexpression of *FUL2* and *FUL3* on the number of spikelets per spike.** (A) Effect of *Ubi::FUL3* on the number of spikelets per spike in $T_1$ plants from two independent transgenic events: #4 (control n = 10, transgene n= 30) and #5 (control n = 9, transgene n= 31). (B) Effect of *Ubi::FUL2* on the number of spikelets per spike in $T_1$ plants from two independent transgenic events: #1 (control n = 4, transgene n= 23) and #6 (control n = 4, transgene n= 11). (C) $F_2$ plants segregating for *VRN1* and *FUL2* in a *vrn2*-null background. (D) $F_2$ plants segregating for *VRN2* and *FUL2* in a *vrn1Vrn1* heterozygous background. (C-D) *P* values correspond to a 2 x 2 factorial ANOVA (N = 35). Error bars are SEM. * = $P < 0.05$, *** = $P < 0.0001$, NS = $P > 0.05$.

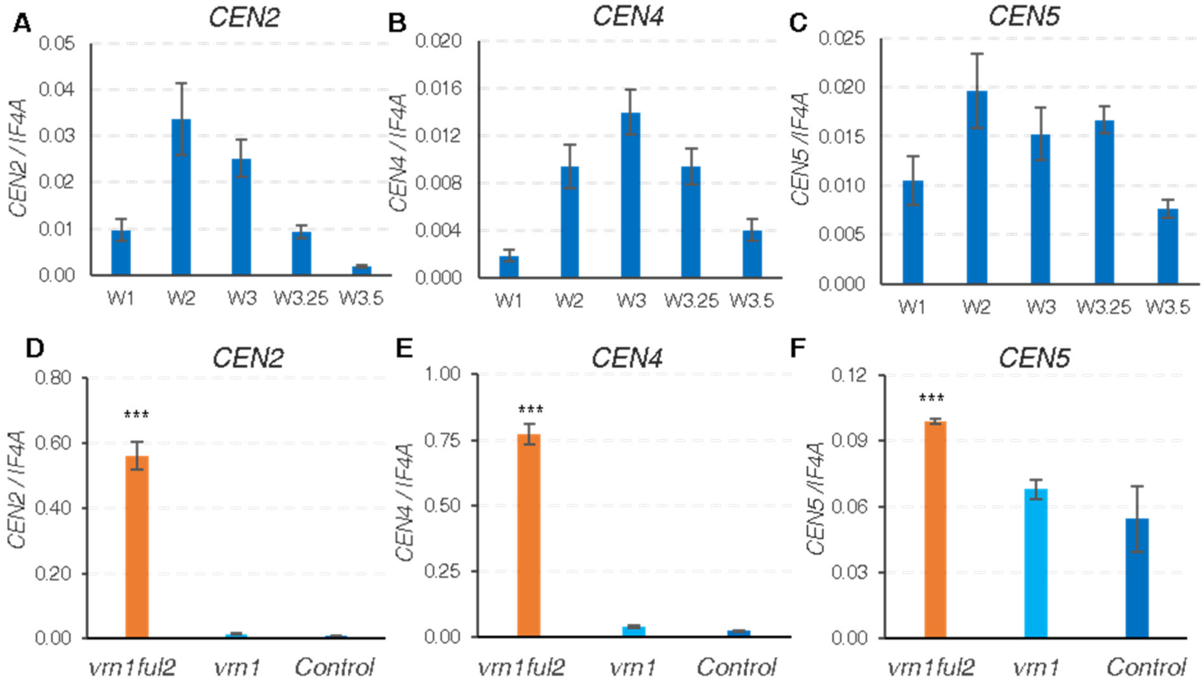


**Fig. S9. Transcript levels of wheat *TFL1/CEN*-like genes *CEN2*, *CEN4* and *CEN5*.** Relative expression levels of (A &D) *CEN2* (TraesCSU01G202000 and TraesCS2B01G310700), (B & E) *CEN4* (TraesCS4A01G409200 and TraesCS4B01G307600), (C & F) *CEN5* (TraesCS5A01G128600 andTraesCS5B01G127600). (A-C) Normal spike development in wild type Kronos *(Vrn1Vrn2*) using the Waddington scale (W1.0= vegetative stage, W2.0 = double ridge stage, W3.0 = glume primordium present, W3.25 = lemma primordium present, W3.5 = and floret primordium present). (D-F) *vrn1ful2*-null, *vrn1*-null, and control at W3.5 (in *vrn2*-null background). Transcript levels were calculated relative to the *INITIATION FACTOR 4A* (*IF4A*) as endogenous control using the ΔCt method. *** = $P < 0.0001$.

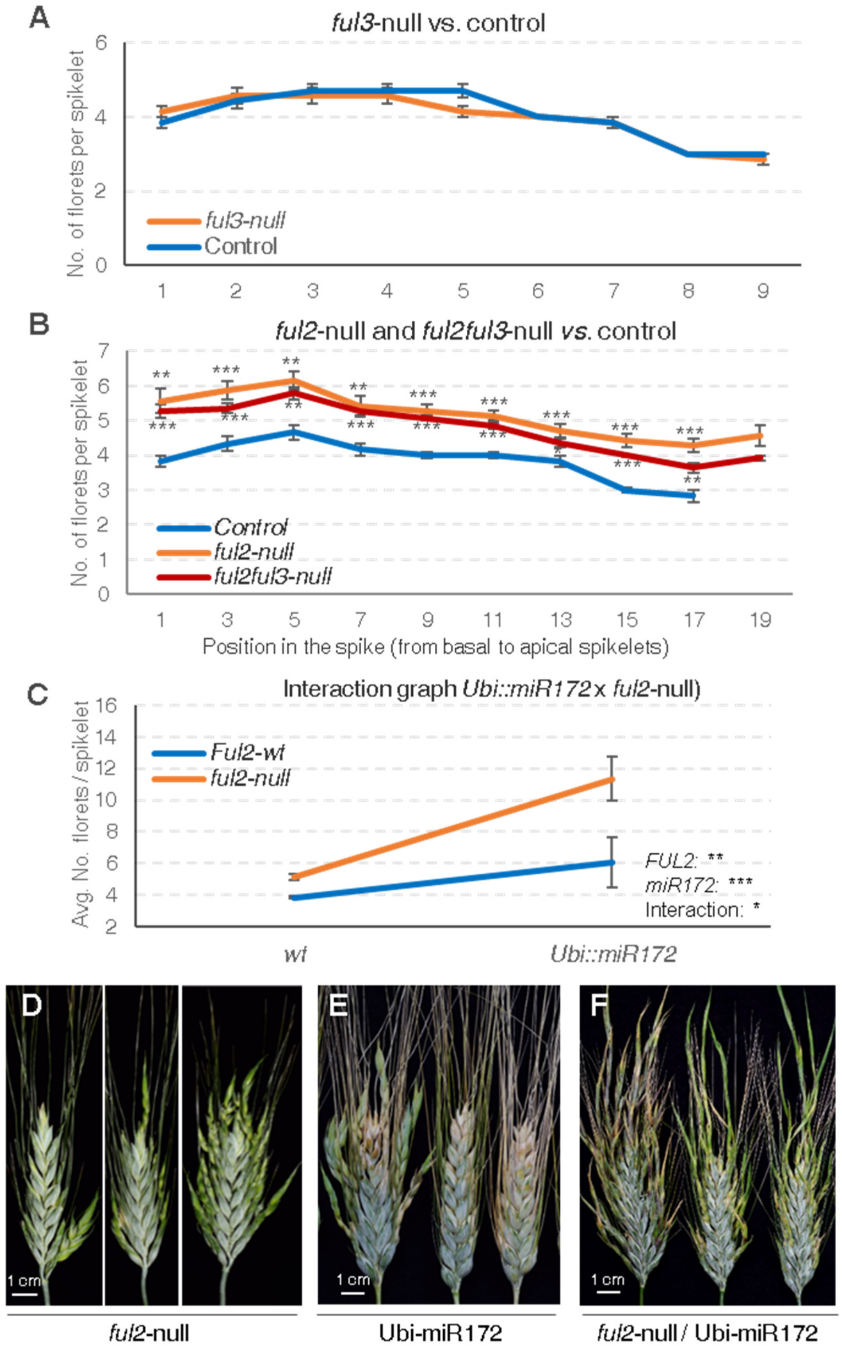


**Fig. S10. Effect of *ful2*-null *ful3*-null and over-expression of *miR172* on the number of florets per spikelet**. (A-B) Distribution of the number of florets per spikelets at different positions in the spike (florets from only one side of seven primary spikes were analyzed). (A) *ful3*-null *vs*. control. (B) *ful2*-null and *ful2ful3*-null *vs*. control (*P* values of each mutant versus the control, by position in the spike). (C) Interaction graph for floret number in $F_2$ plants from the cross *Ubi::miR172* x *ful2*-null. (D-F) Representative spikes showing heterogeneity in the distribution of spikelets with extra florets in (D) *ful2*-null, (E) *Ubi::miR172* and (F) *ful2*-null + *Ubi::miR172*. ** = $P < 0.01$, *** = $P < 0.001$, error bars are SEM.
